# Membrane Env liposomes for immunization with HIV spikes

**DOI:** 10.1101/2020.11.29.403204

**Authors:** Daniel P. Leaman, Armando Stano, Yajing Chen, Lei Zhang, Michael B. Zwick

## Abstract

A key goal in HIV vaccine design remains to elicit broadly neutralizing antibodies (bnAbs) against the membrane-embedded envelope glycoprotein spike (mEnv). However, mEnv has lagged behind engineered soluble Envs in vaccine development due to low expression yields and the presence of extraneous proteins on particles. Here, we describe a mEnv vaccine platform that requires no extra proteins or protein engineering. MEnv trimers were fixed, purified and combined with liposomes in mild detergent. On removal of detergent, mEnvs were observed embedded in particles, designated <u>mE</u>nv <u>l</u>iposomes (MELs), which were recognized by HIV bnAbs but not non-nAbs. Following sequential immunization in rabbits, MEL antisera neutralized select tier 2 HIV isolates. Variations between the Env immunogens, including a missing N-glycosylation site at position 197 near the CD4 binding site, provide insights into the specificities elicited and possible ways to improve immunogens. MELs can facilitate vaccine design to elicit HIV bnAbs using biochemically defined and multimerized mEnv.

## Introduction

HIV/AIDS afflicts more than 38 million people worldwide and is currently without a practical cure (UNAIDS, 2020). Modest efficacy of a vaccine against HIV infection has been correlated to the activity of certain non-neutralizing antibodies in humans (Rerks-Ngarm et al., 2009; Tomaras & Plotkin, 2017). However, to effectively curb the pandemic that involves a high diversity of circulating viral isolates, a vaccine most likely will need to reproducibly elicit HIV broadly neutralizing antibodies (bnAbs), something which described vaccines have failed to do (Bjorkman, 2020; Haynes, Burton, & Mascola, 2019).

HIV bnAbs must recognize the envelope glycoprotein trimeric spike in its membrane-embedded state (mEnv). Many early vaccines consisted of glycoprotein (gp) subunits of Env, *e.g*. gp120 outer subunit, gp160 precursor, or soluble gp140s that were uncleaved by furin and devoid of the transmembrane (TM) domain and C-terminal tail (CTT) of subunit gp41 (Spearman, 2006). These subunit vaccines elicited antibodies with limited neutralization activity that targeted variable epitopes (Gilbert et al., 2010; Mascola et al., 1996; Spearman et al., 2011; Yang, Wyatt, & Sodroski, 2001), but they lacked well-ordered trimeric structure and were missing important bnAb epitopes (Ringe et al., 2013).

More recently, soluble (s)Env gp140s have been engineered with trimer stabilizing mutations, *e.g*. SOSIP (Sanders et al., 2013; Sanders et al., 2002), UFO (Kong et al., 2016), and NFL (Sharma et al., 2015), that reasonably mimic the structure of mEnv, recapitulate quaternary bnAb epitopes and occlude many non-nAb epitopes (Julien et al., 2013; Lee, Ozorowski, & Ward, 2016; Stadtmueller et al., 2018). Immunization with these sEnvs has elicited nAbs against relatively resistant (tier 2) primary viruses that often target variable loops or gaps in Env’s glycan shield (Crooks et al., 2015; Hessell et al., 2016; McCoy et al., 2016; Sanders et al., 2015). NAb responses have been enhanced in some cases in which sEnv immunogens were chemically crosslinked (Dubrovskaya et al., 2019; Leaman, Lee, Ward, & Zwick, 2015; Schiffner et al., 2018), multimerized on nanoparticles or liposomes (Bale et al., 2017; McGuire et al., 2016; Morris et al., 2017), and/or engineered with certain N-linked glycosylation site mutations. A sequential prime-boost regimen using such approaches has recently elicited sporadic titers of bnAb (Dubrovskaya et al., 2019).

Immunization with sEnvs often elicits disproportionately high titers of non-nAbs to the truncated base (Bale et al., 2017; Hu et al., 2015) while key epitopes of bnAbs 2F5, 4E10 and 10E8 against the membrane-proximal external region (MPER) are either disrupted or missing (Krebs et al., 2019; Zhang et al., 2019). Differences in conformational dynamics between sEnv and native mEnv have also been reported (Lu et al., 2019). Virus-like particles (VLPs) displaying mEnv have been used as immunogens but may elicit off-target or distracting antibody responses to Env debris or extraneous proteins from the virus and the host cell, which may also be autoreactive (Cantin, Methot, & Tremblay, 2005; Crooks et al., 2007; Leaman et al., 2015; Poon, Hsu, Gudeman, Chen, & Grovit-Ferbas, 2005). Genetic vaccines such as mRNA vaccines have gained recent attention as a promising platform (Pardi, Hogan, Porter, & Weissman, 2018), but do not allow control over the conformation, density, as well as quality of the protein produced and does not allow crosslinking for added stability. Hence, a suitable mEnv vaccine alternative is desirable that addresses the above concerns, to diversify approaches and enable new hypotheses.

Here, we developed a vaccine platform that displays multivalent, well-ordered mEnv spikes on liposomes, termed MELs. MEnv was purified readily from cell lines (Stano et al., 2017) and assembled into liposomes with MPER exposed and CTT buried. Notably, MELs were devoid of extraneous proteins and displayed stable, crosslinked mEnv trimers in a multivalent array. In an immunization experiment, MELs elicited antibodies that neutralized select tier 2 isolates of HIV. Hence, MELs should be a useful platform for rational HIV vaccine design involving mEnv.

## Results

### Generation of fixed mEnv spikes

We previously described an HIV Env nanoparticle immunogen in which mEnv was captured on antibody-coated nanobeads (Leaman et al., 2015). In that approach, the capture antibody was an undesirable vaccine component, and mEnv yields were low from the transient transfection of cells. Here, we took advantage of a recently described stable cell line strategy that increased mEnv yield by more than 10-fold (Stano et al., 2017). The mEnv of this cell line is a variant of the clade B isolate ADA, termed Comb-mut (ADA.CM), which was previously selected for high trimer stability (Leaman & Zwick, 2013). We also generated two new mEnv cell lines, JRFL.TD15 and CH505.N197D, to produce mutant Envs of a clade B isolate and a clade C transmitted/founder (T/F) isolate, respectively, as will be described further below.

MEnv-expressing cells were fixed in glutaraldehyde (GA), a clinically approved chemical crosslinker shown previously to have helped in eliciting HIV nAbs (Schiffner et al., 2018; Soldemo et al., 2017). Fixed cells were solubilized in the detergent n-dodecyl-β-D-maltoside (DDM) and mEnv was affinity purified using the trimer specific antibody PGT151. Size exclusion chromatography of purified mEnv revealed two major peaks, the first being aggregated mEnv, while the second was trimeric mEnv that was used for subsequent biophysical studies (**Supp. Fig. 1A-B**). Purified mEnvs ADA.CM, CH505.N197D and JRFL.TD15 were verified to be trimeric using blue native (BN)-PAGE and denaturing SDS-PAGE (**Fig. 1A** and **Supp. Fig. 1B**). The trimers were recognized by several bnAbs to gp120 and gp41, including those to quaternary epitopes, and showed minimal binding by non-nAbs (**Fig. 1B**). These data show the mEnvs are stable, trimeric and have native-like antigenicity. Of note, mEnvs were produced in yields sufficient for immunizations, *i.e*., ~0.5 mg of pure mEnv per liter of culture medium.

**Figure 1.**
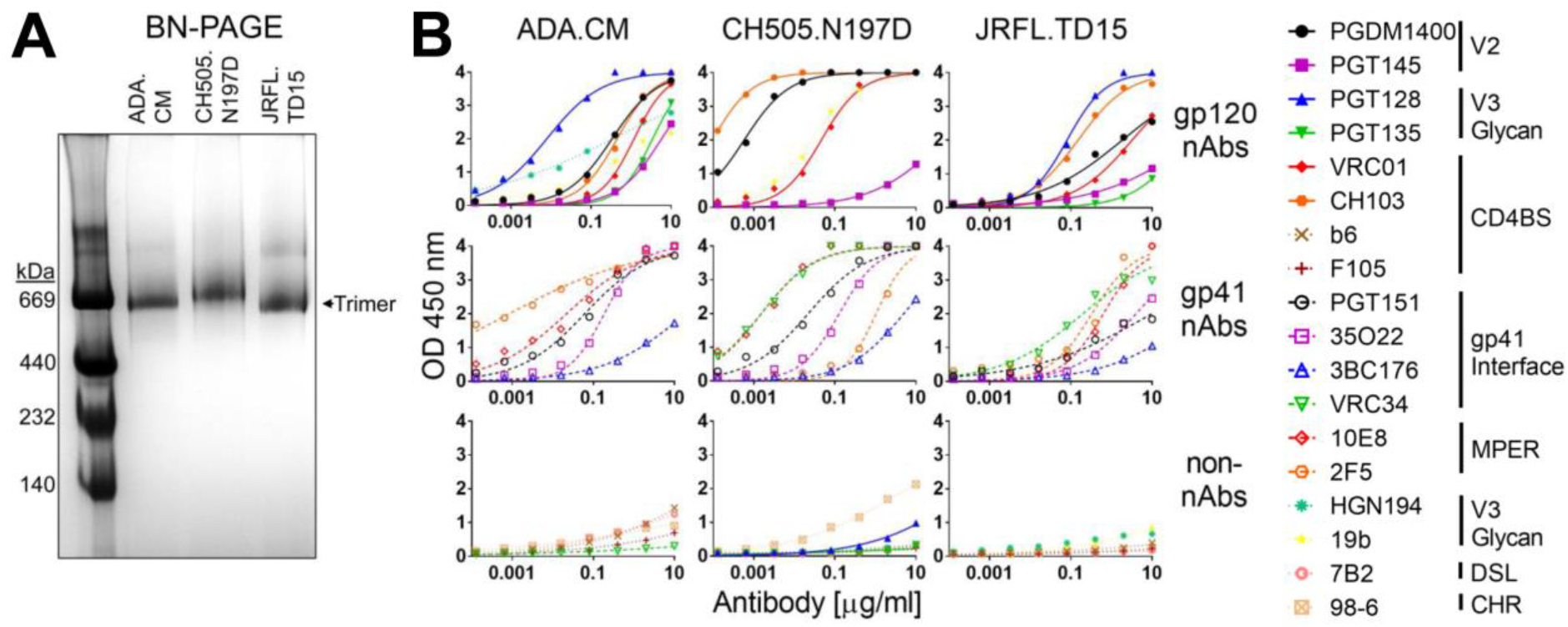
Purified mEnvs are trimeric and display native antigenicity. (**A**) MEnv (5 μg) was analyzed using BN-PAGE stained with Coomassie blue. (**B**) Purified mEnv was captured using lectin from *Galanthus nivalis* (GNL) and probed in an ELISA using a panel of antibodies. Antibodies are classified as nAbs if they neutralize cognate virus with an IC_50_ <50 μg/ml.

### Incorporation of mEnv into liposomes

To see if mEnv could incorporate into liposomes, we adopted an approach described for making proteoliposomes (**Fig. 2A**) (Geertsma, Nik Mahmood, Schuurman-Wolters, & Poolman, 2008; Seddon, Curnow, & Booth, 2004). Thus, naked liposomes (70% POPC, 30% cholesterol), that had been extruded through a membrane with a 100 nm pore diameter, were treated with DDM at a concentration predetermined to saturate - but not fully solubilize - the lipid bilayer. Purified mEnv was added, and then the detergent was removed by adsorption using polystyrene Bio-beads (Rigaud et al., 1997). The mean particle size of both naked liposomes and the mEnv-liposome mixture was ~140 nm as determined by nanoparticle tracking analysis (NTA; **Fig. 2B**). Notably, negative stain electron microscopy (EM) revealed prominent, outward protruding spikes on the liposomes, which we termed membrane-Env liposomes, or MELs (**Fig. 2C**). We probed the antigenicity of the MELs using a panel of antibodies in an ELISA. As expected, the MELs were bound efficiently by most bnAbs, while binding by non-nAbs was minimal (**Fig. 2D**). We also probed MELs using the anti-CTT antibody, Chessie8 (Steckbeck, Sun, Sturgeon, & Montelaro, 2013), and found that it did not bind the MELs but did bind to detergent-solubilized mEnvs, which is in agreement with the EM images showing that MELs were decorated with embedded spikes pointing outwards. We conclude that mEnv incorporated into the bilayer of MELs in the correct orientation with the ectodomain and MPER accessible to bnAbs and the CTT buried within the liposome.

**Figure 2.**
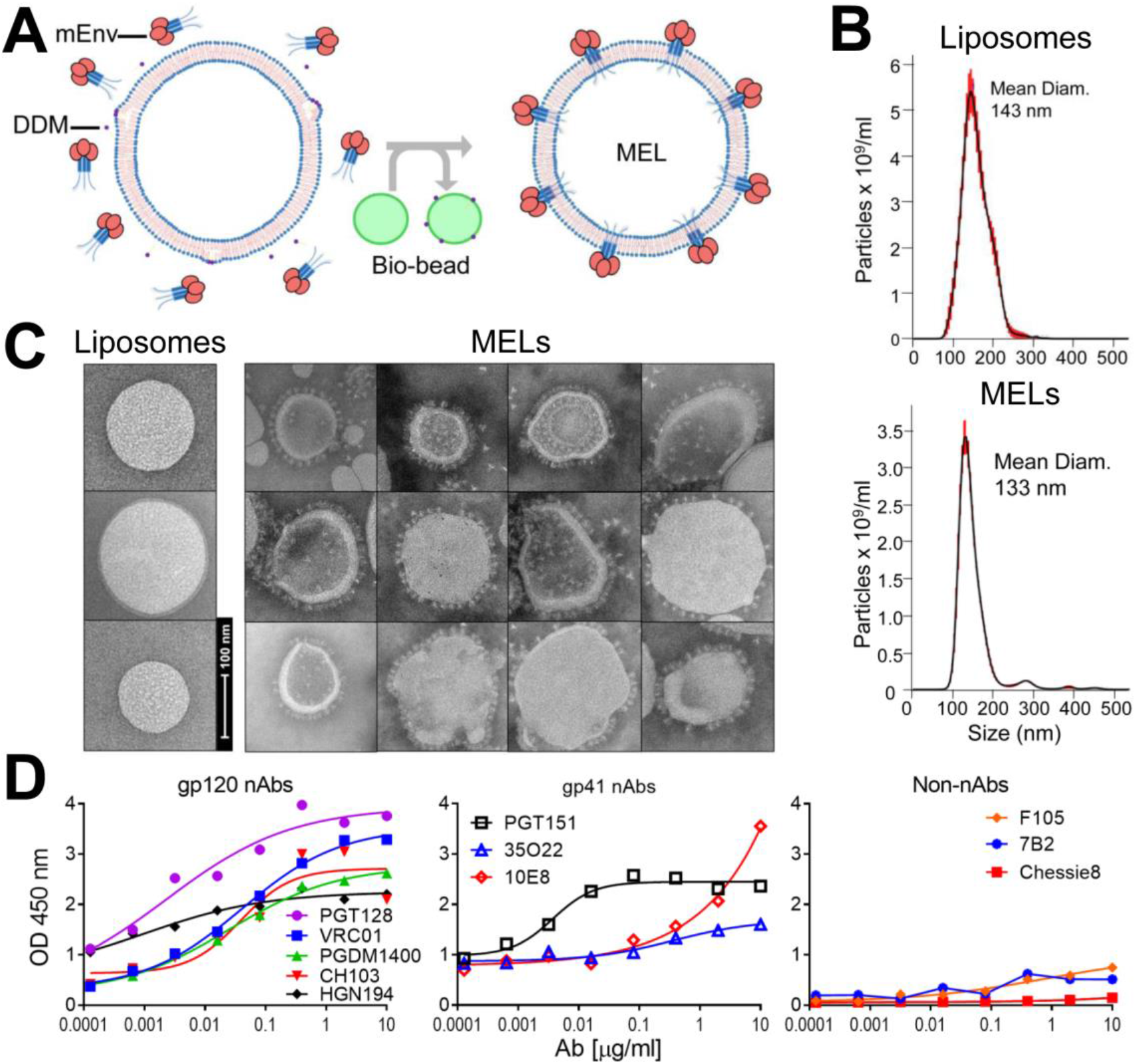
Production, properties and antigenicity of mEnv liposomes (MELs). (**A**) Naked liposomes were treated with mild detergent DDM that destabilizes the membrane bilayer and then combined with purified mEnv trimers (left). Detergent was depleted from the liposome-mEnv mixture by repeated incubations with bio-beads, which yields MELs decorated with an array of spikes (right). (**B**) Nanoparticle tracking analysis (NTA) reveals liposomes are monodispersed with a mean diameter of 143 nm; MELs with mEnv remain monodispersed with a mean diameter of 133 nm. (**C**) Negative stain EM shows MELs embedded with mEnv trimeric spikes. (**D**) ELISA data shows bnAbs bind to MELs but non-nAbs and anti-CTT antibody, Chessie8, do not. MELs were labeled with biotin-DOPE and captured on streptavidin coated microwells.

The stability of MELs was analyzed over time at physiological temperature using negative stain EM and a capture ELISA. MELs appeared to retain the embedded spikes and overall appearance when visualized at 96 h by negative stain EM (**Supp. Fig. 2A**). Additionally, after 7 days at 37°C MELs were recognized efficiently by nAbs to the CD4 binding site (CD4BS), N332 glycan supersite, V3, and MPER (**Supp. Fig. 2B**), while binding by three quaternary bnAbs, PGDM1400, PGT151, and 35O22, modestly decreased. Importantly, binding by non-nAbs or anti-CTT antibody to MELs did not increase after the incubation. We surmised trimeric mEnv and MELs were likely stable enough to elicit relevant humoral responses in animals.

### MELs elicit HIV neutralizing antibodies

Next, we explored the immunogenicity of MELs. We chose to immunize with MELs in Alum plus CpG using New Zealand White (NZW) rabbits, as we had used these adjuvants for a prior rabbit study in which sporadic tier 2 autologous nAb responses were elicited (Leaman et al., 2015). We note that although CpG ODN 1826 reportedly blocked binding of V2 bnAbs to SOSIP gp140 (Ozorowski et al., 2018), we saw no effect of CpG ODN 2007 on the reactivity of V2 or other bnAbs to mEnvs in an ELISA (**Supp. Fig. 3**). Hence, rabbits were inoculated with ADA.CM MELs every six weeks for four immunizations (**Fig. 3A**).

**Figure 3.**
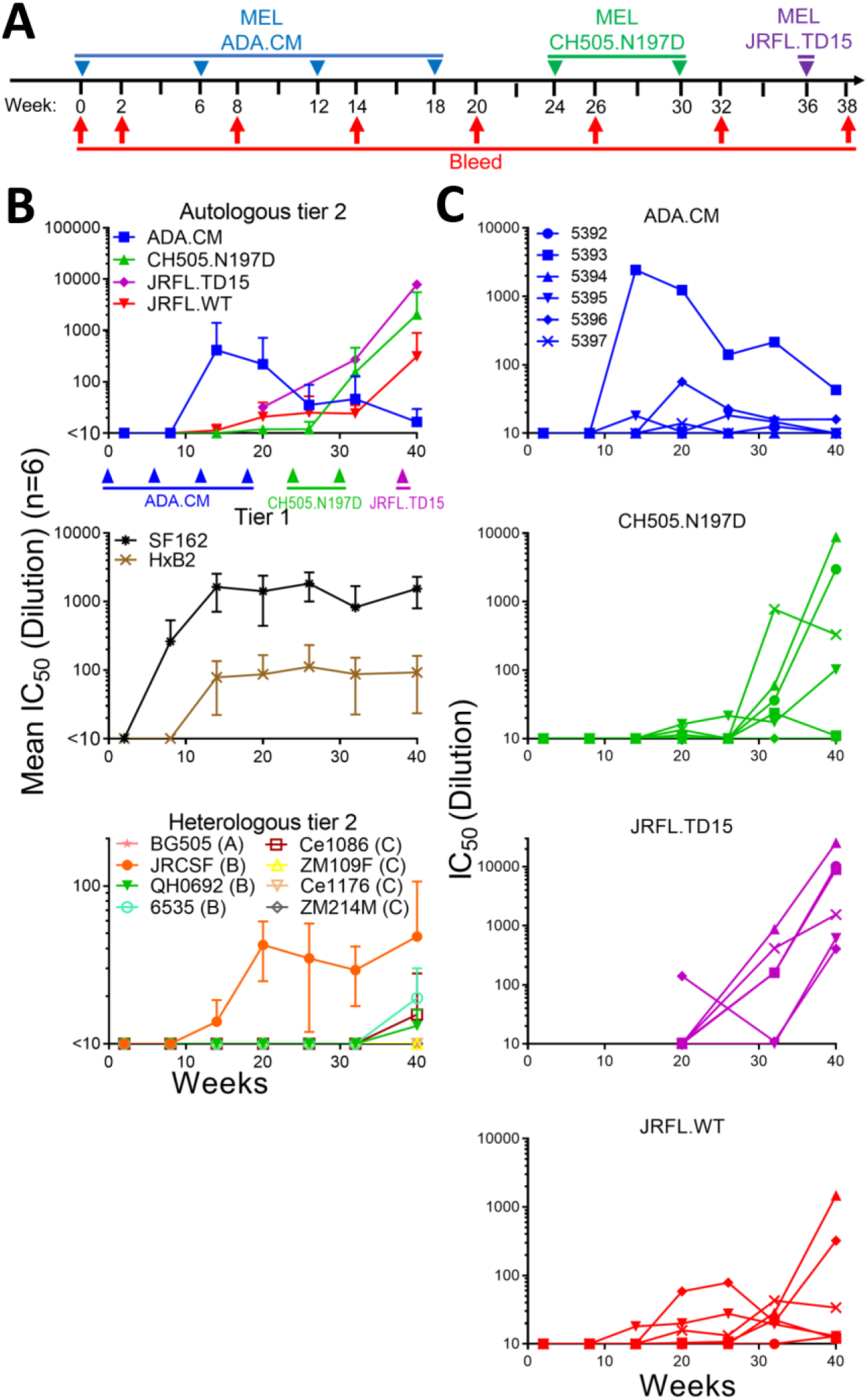
Immunization with MELs in rabbits elicited HIV nAbs. (**A**) Schedule of immunization of NZW rabbits with MELs. Rabbits were immunized with MELs seven times, *i.e*., ADA.CM (4 times), CH505.N197D (2 times), and JRFL.TD15 (1 time), six weeks apart. (**B**) Kinetics of neutralization of autologous and heterologous tier 1 and tier 2 isolates by MEL immune sera. Mean IC_50_s of six rabbit sera from each bleed against the three immunizing or autologous isolates and JRFL.WT (top), heterologous tier 1 isolates (middle), and a panel of heterologous tier 2 isolates (bottom). (**C**) Kinetics of neutralization of immunizing virus strains by individual rabbit sera.

Sera from ADA.CM immunized rabbits were tested for HIV neutralizing activity in a standardized assay using TZM-bl target cells. Tier 1a strains SF162 and MW965 were neutralized with mean IC_50_s >1:1,000, while neutralization of tier 1b strain HxB2 was modest but consistent with a mean IC_50_ of about 1:100 (**Fig. 3B** and **Supp. Fig. 4**). Sera from four of six rabbits showed autologous neutralization against ADA.CM, with rabbit serum 5393 reaching an IC_50_ of >1:1,200 (**Fig. 3B** and **C**). Heterologous primary isolates were also neutralized by the sera, including the clade B isolates JRCSF (mean IC_50_ of 1:42 for six responders) and JRFL (mean IC_50_ of 1:31 for four responders), while a clade C mutant virus CH505.N197D was also neutralized (mean IC_50_ of 1:14 for four responders).

Encouraged by the MEL ADA.CM immunogenicity results, we chose to boost the animals using a phylogenetically distant Env, anticipating that such a strategy might broaden serum neutralizing activity if sera could already weakly neutralize the boosting mEnv. For the first boost, we chose CH505.N197D since 4 out of 6 rabbit sera showed detectable neutralization of this mEnv and a stable cell line of it had already been prepared. CH0505 is a transmitted/founder (TF) virus from a donor that developed bnAbs to the CD4BS (Liao et al., 2013), and CH0505.N197D is an N-glycosylation site mutant that showed increased sensitivity to CD4BS bnAbs without increased binding by non-neutralizing CD4BS antibodies b6 and F105 (**Fig. 1B** and **7B**). Following two boosts with CH505.N197D MELs, nAb titers increased against CH505.N197D virus, albeit less so toward CH505 wild-type (WT) with an intact N197 N-glycosylation site (**Supp. Fig. 4**). Notably, rabbit 5397 neutralized CH505.N197D with an IC_50_ of ~1:1,000. Meanwhile, although serum neutralization of ADA.CM decreased, neutralization against ADA and JRCSF was unchanged, and nAb titers against the heterologous isolate JRFL became more consistent (**Fig. 3B, C** and **Supp. Fig. 4**).

Because boosting with CH505.N197D improved heterologous neutralization, we chose to boost the animals once more. We had generated a mEnv cell line previously, JRFL.TD15, which incorporates “TD15” mutations shown to stabilize soluble JRFL gp140 trimers (Guenaga et al., 2015). The functional stability of JRFL.TD15 mEnv, *i.e*., the temperature (T90) at which an Env pseudovirus loses 90% infectivity in an hour (Agrawal et al., 2011), was indeed found to be 3°C higher than JRFL.WT (**Supp. Fig. 5A**). Of note, JRFL.TD15 showed increased sensitivity to V2 and V3 antibodies as compared to JRFL.WT (**Supp. Fig. 5B**). JRFL.TD15 virus was also more sensitive than JRFL.WT to 4 of 6 rabbit sera following boosting with CH505.N197D MELs (**Supp. Fig. 4**). The boost with JRFL.TD15 MELs strongly increased nAb titers against JRFL.TD15 virus in all animals, up to >1:25,000 (**Supp. Fig. 4** and **5C**). Neutralization titers against JRFL.WT were lower but had increased in 4 of 6 animals, with 50-fold and 15-fold increases in animals 5394 and 5396, respectively. Neutralization also increased against CH505.N197D with several sera, up to 1:5700 (**Supp. Fig. 4**). Heterologous neutralization - albeit sporadic and weak - increased against JRCSF, and, notably, could now be observed against the clade Bs QH0692 and 6535, and the clade C isolate Ce1086.

### Antibody binding specificities in MEL immune sera

MEL ADA.CM immune sera were probed in an ELISA and found to contain high antibody binding titers against trimeric ADA.CM and monomeric gp120 in ELISA, with lower titers to recombinant gp41 (**Fig. 4A**). Serum antibodies also bound to the full MPER peptide (a.a. 654-683) containing the 2F5 and 4E10/10E8 epitopes, but not to the C-terminal MPER peptide (a.a. 670-683), suggesting that elicited MPER antibodies targeted mostly N-terminal epitopes in the MPER (**Supp. Fig. 6A**). Serum antibodies bound to V3 peptides of gp120 and the immunodominant Kennedy epitope region in the gp41 CTT. The latter result indicates the CTT was exposed to B cells, perhaps from disrupted MELs. Antibody titers to the FP were detectable but low, and no binding was observed to the gp41 disulfide loop that is occluded in mEnv trimer structures (Julien et al., 2013; Sanders et al., 2013; Stano et al., 2017).

**Figure 4.**
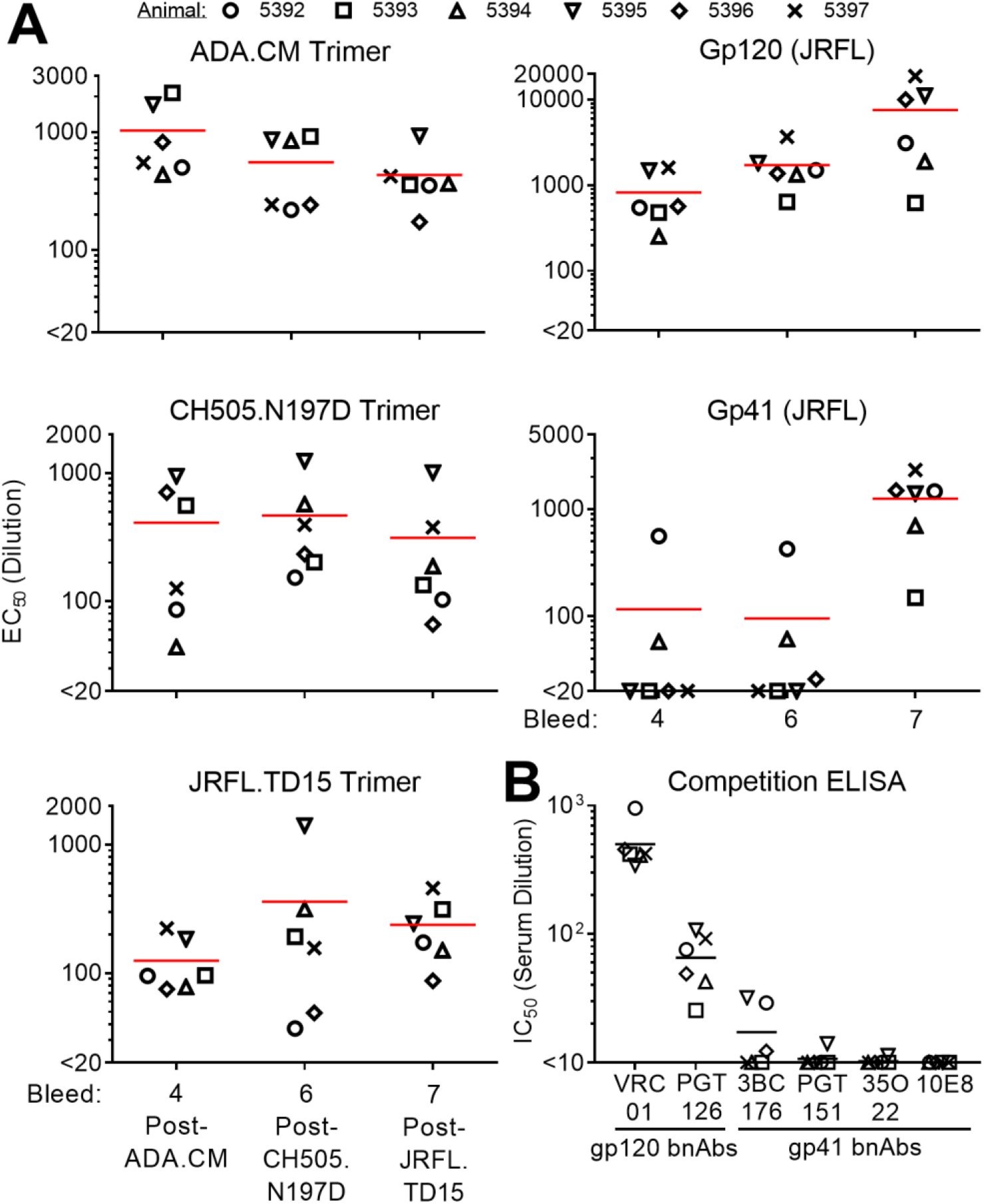
Antibody binding specificities in MEL antisera revealed by ELISA. (**A**) Sera from bleeds 4 (post-ADA.CM), 6 (post-CH505.N197D), and 7 (post-JRFL.TD15) were tested for binding to each of the three immunizing mEnvs, as well as recombinant gp120 (JRCSF) and gp41 (JRFL). (**B**) Sera from the final bleed were tested for the ability to block biotinylated bnAb binding to JRFL.TD15 trimer. The biotinylated bnAbs include those against the CD4BS (VRC01), N332 glycan supersite (PGT126), gp120-gp41 interface (3BC176, PGT151, and 35O22) and the MPER (10E8).

The immune sera following boosting with CH505.N197D and JRFL.TD15 MELs showed diminished titers in ELISAs against gp41 epitopes MPER, FP, and CTT, while titers to monomeric gp120 rose about 10-fold (**Fig. 4A** and **Supp. Fig. 6A**). Of possible relevance, some N-terminal MPER, FP, and CTT residues are mismatched between ADA.CM, CH505.N197D, and JRFL.TD15 (**Supp. Fig. 6B**). Titers against V3 peptide decreased to undetectable levels from boosting with CH505.N197D, perhaps due to limited exposure of the V3 crown (**Fig. 1B**) and sequence differences in V3 between CH505.N197D and ADA.CM. V3 antibody titers rebounded following the boost with JRFL.TD15 that is modestly sensitive to V3 crown antibodies (**Supp. Fig. 6A**).

Immune sera from the final bleed were also tested in a competition ELISA for the ability to block bnAbs from binding to JRFL.TD15 mEnv. All six sera robustly blocked binding by VRC01 and PGT126 to the CD4BS and N332 glycan supersite, respectively. (**Fig. 4B**). Two sera, which were shown to recognize a CHR peptide (**Supp. Fig. 6**), weakly blocked binding of bnAb 3BC176 to the gp120-gp41 interface (**Fig. 4B**). Altogether, serum specificities to Env seem to be diverse, and often overlap with the CD4BS or N332 supersite, and less often with the gp41 epitopes.

### Neutralization specificities in MEL immune sera

Next, we determined the HIV nAb specificities elicited by MEL immunization. We found the most potent neutralizing serum against ADA.CM, 5393, did not neutralize the parental isolate, ADA. Of the seven mutations that differentiate ADA.CM from ADA, we found the V1 alteration (N139/I140 deletion, N142S) was sufficient to make ADA as sensitive to 5393 as ADA.CM and hence is the likely target of nAbs in this serum.

Removal of consensus N-linked glycosylation sites on Env (*i.e*., glycan holes) naturally exposes the underlying protein surface and has been associated with eliciting nAbs against autologous HIV (Crooks et al., 2015; McCoy et al., 2016; Voss et al., 2017; Zhou et al., 2017). ADA.CM lacks two relatively conserved N-linked glycosylation sites at N289 and N230, so we asked whether the sera could neutralize ADA.CM mutants with N-glycosylation sites restored at these sites (K289N and D230N/K232T), as well as with other N-glycosylation site knockouts near this gp120-gp41 interface (N88A, N241S, N611D and N637K). The N-glycosylation site mutations did not affect neutralization by most sera, but the four knockout mutations did significantly increase neutralization by serum 5396, suggesting that some nAbs in this serum target the gp120-gp41 interface (**Fig. 5B**).

**Figure 5.**
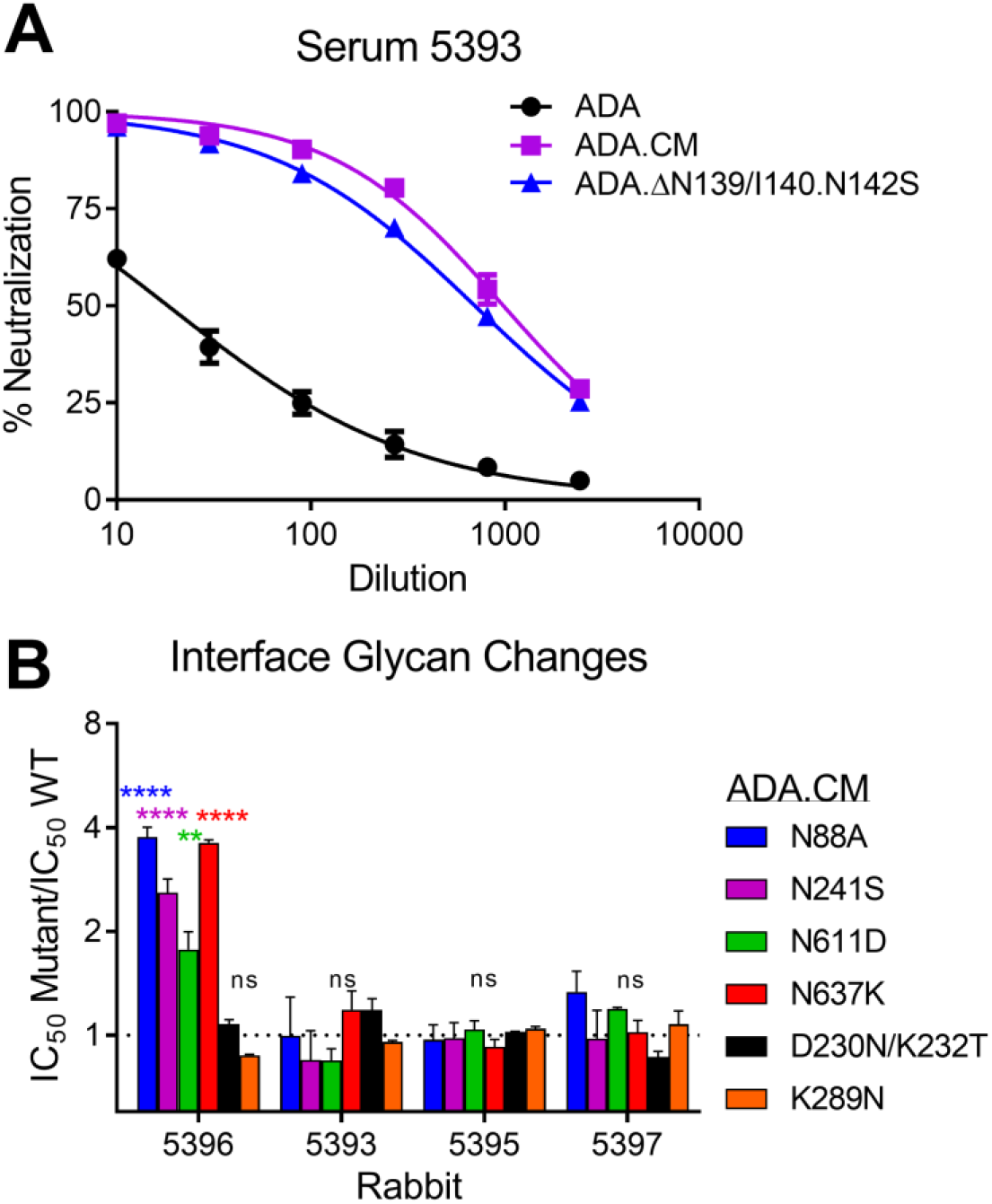
ADA.CM neutralization activities in sera 5393 and 5396 target V1 and the gp120-gp41 interface. (**A**) Serum 5393 from Bleed 3, following immunization with ADA.CM, was tested for neutralization of ADA.CM, parental ADA, and an ADA mutant containing a N139/I140 deletion and N142S substitution in V1 of gp120. (**B**) Four rabbit sera from Bleed 4 that neutralized ADA.CM with an IC_50_ >10 were tested in neutralization assays against ADA.CM with glycan knockout mutations (N88A, N241S, N611D and N637K) or glycan hole filling mutations (D230N/K232T and K289N) near the gp120-gp41 interface.

We used V3 crown and MPER linear peptides as “dump-in” reagents to see if they could block HIV serum neutralization. As anticipated, V3 peptide blocked neutralization by anti-V3 antibody F425-B4e8 and blocked most serum neutralization of the Tier 1a SF162 strain (**Fig. 6A-B**). Neutralization of heterologous tier 2 isolate Ce1086 was also abrogated by addition of V3 peptide, as was neutralization of JRCSF but only in two of three animals. V3 peptide competition did not affect the potent neutralization of JRFL.WT and CH505.N197D, and only decreased neutralization of JRFL.TD15 and heterologous Tier 2 isolate QH0692 about two-fold. An MPER peptide did not block neutralization with any sera or isolate tested (data not shown).

**Figure 6.**
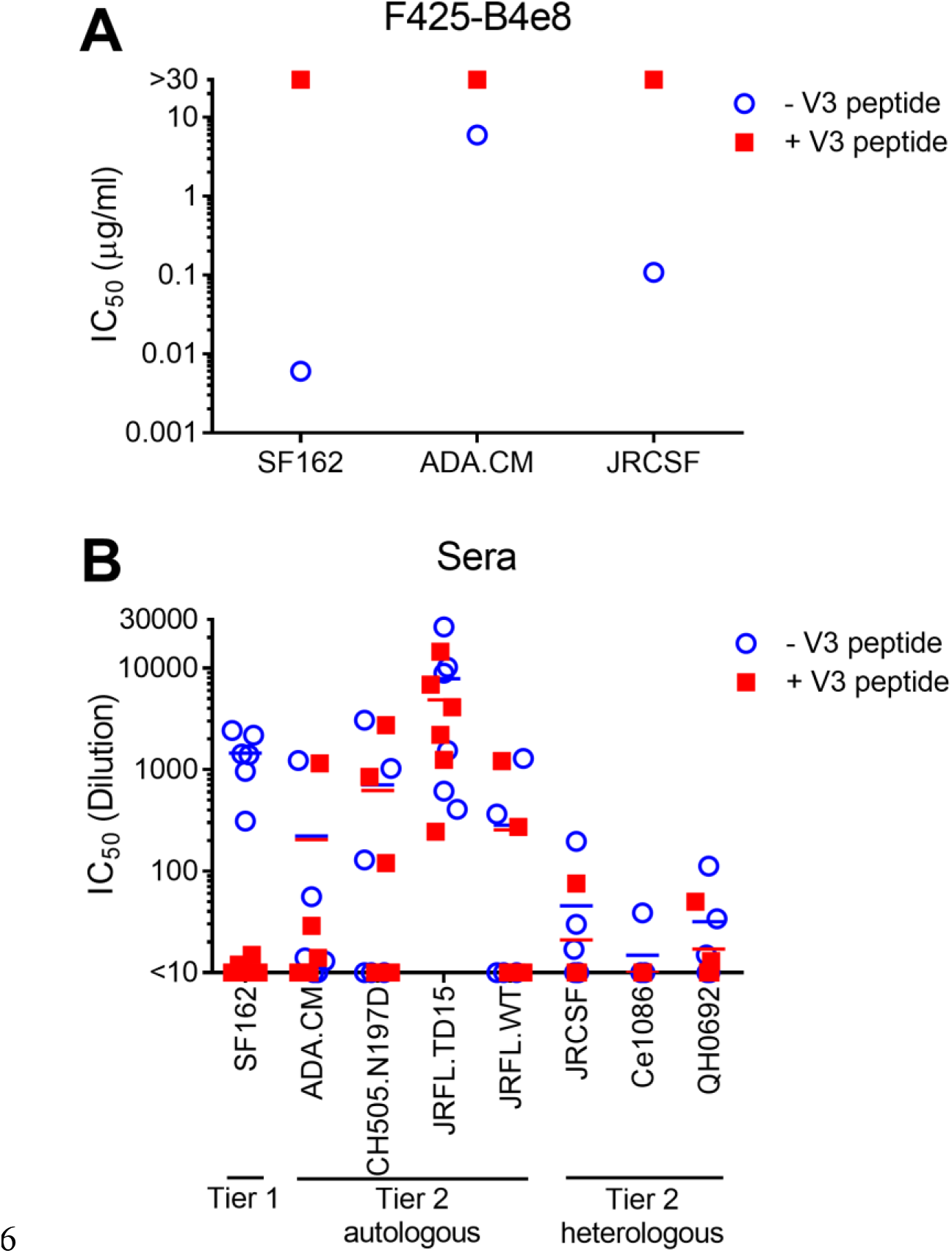
Serum neutralization of autologous isolates and some heterologous Tier 2 isolates cannot be blocked by V3 peptide. Neutralization by (**A**) monoclonal antibody F425-B4e8 that binds to the V3 crown, and (**B**) sera from the final bleed, was tested in a neutralization assay against HIV isolates in the presence or absence of V3 peptide (JRFL sequence: NTRLSIHIGPGRAFYTTGEIIGDI).

To further assess the neutralization discrepancy between JRFL.TD15 and JRFL.WT, we generated R308H and WT-V2 mutants of JRFL.TD15, that revert V3 and V2, respectively, to that of JRFL.WT. On average R308H decreased the IC_50_ of the sera 8-fold, ranging from no change (serum 5396) to a 24-fold decrease (serum 5395), so it accounted for some but not all of the difference in sensitivity between JRFL.TD15 and JRFL.WT (**Supp. Fig. 5C**).

JRFL.TD15.WTV2, on the other hand, decreased the neutralization of JRFL.TD15 back to JRFL.WT levels for 4 of the 6 sera, suggesting that most JRFL.TD15 nAbs that do not neutralize JRFL.WT target V2.

V2 nAbs have been reportedly elicited by soluble CH505 immunogens (Saunders et al., 2017). Hence, we generated an N160A mutant of CH505.N197D that knocks out neutralization by V2 bnAbs (Walker et al., 2009). Removing the N160 glycosylation site did not affect serum neutralization of CH505.N197D, suggesting the elicited nAbs do not target this conserved region of V2 (**Fig. 7C**).

**Figure 7.**
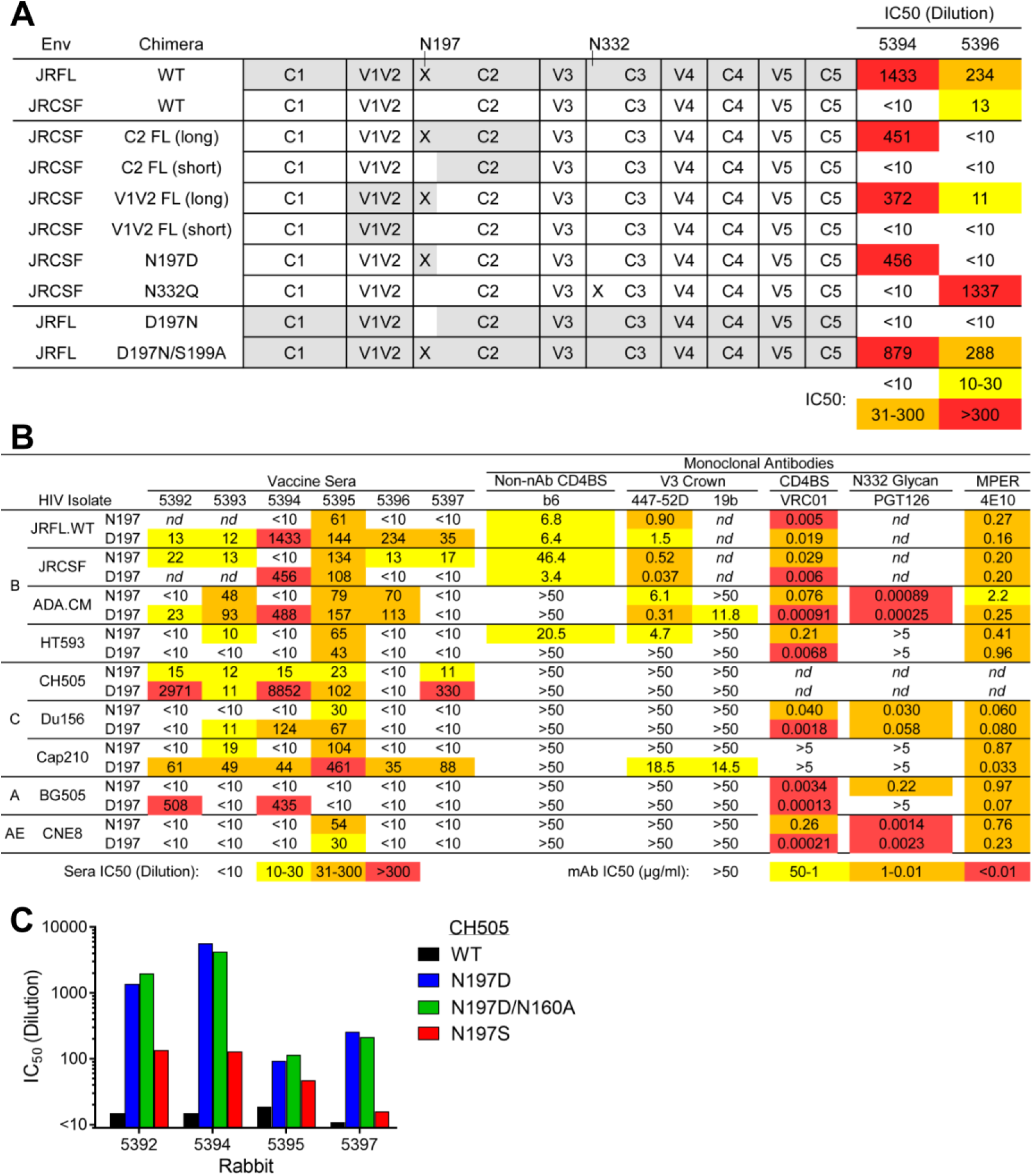
MELs elicited potent nAbs to a glycan hole at position 197. (**A**) Neutralization by sera 5394 and 5396 against a panel of domain swapped HIV Env chimeras between JRFL and the more resistant JRCSF, as well as against N-glycosylation site mutants N197 and N332 of JRCSF. (**B**) N197D mutants of diverse HIV isolates were tested for neutralization by sera from the final bleed post-sequential MEL immunization (left) as well as by a panel of bnAbs and non-nAbs (right). JRFL and JRCSF and their N197 mutants were tested for neutralization by V3 antibody F425-B4e8 and MPER antibody 2F5 rather than 447-52D and 4E10, respectively. (**C**) Rabbit sera from the final bleed were tested for neutralization against CH505.WT, CH505.N197D, CH505.N197D.N160A that knocks out V2 bnAb neutralization, and CH505.N197S.

Envs CH505.N197D and JRFL.TD15 both lack N-linked glycosylation sites at position 197 near the CD4BS, so we anticipated some nAbs to these viruses may target the common glycan hole. We made use of the fact that sera from two animals, 5394 and 5396, neutralized JRFL but not the tier 2 isolate JRCSF that has the N197 glycosylation site. Domain substituted chimeras of JRFL and JRCSF were therefore tested for neutralization by these sera. Serum 5394 neutralized JRCSF.N197D and JRCSF chimeras engrafted with JRFL V2 or C2 domains that also introduced D197, while 5396 did not (**Fig. 7A**). However, an N-linked glycosylation site knock-in mutation D197N in JRFL abrogated neutralization by both sera. Of note, serum 5396 potently neutralized JRCSF N332Q, but serum 5394 did not. Overall then it seems each serum has distinct nAb specificities against the 197 glycan hole on JRFL.

To understand antibody specificities to the N197 glycan hole in more detail, we made N197D mutants of isolates ADA.CM, HT593, Du156, CAP210, BG505, and CNE8, in addition to mutants of CH505, JRCSF, and JRFL described above. Sera 5396 and 5397 only neutralized JRFL and CH505.N197D, respectively (**Fig. 7B**). However, serum 5394 could neutralize Du156, BG505, ADA.CM, Cap210, CH505, JRCSF, and JRFL isolates with D197, for a total of 7 out of 10 N197 glycan-deleted isolates. Serum 5392 neutralized BG505.N197D (IC_50_=1:500) as well as CH505.N197D, thus revealing a modest breadth against N197D mutants.

Serum neutralization was also compared using CH505.N197D and CH505.N197S. The N197S mutant of CH505 was less sensitive than N197D to all tested sera but was more sensitive than CH505.WT (**Fig. 7C**). These data suggest some nAbs elicited to the CH505 N197 glycan hole may depend in part on the Asp side chain at position 197.

The V5 loop of gp120 plays a role in the epitopes of many CD4BS bnAbs (Fera et al., 2014; Schommers et al., 2020; Umotoy et al., 2019). We therefore generated V5 mutants of JRFL and CH505 to test whether serum nAbs targeting the N197D glycan hole have such features in common with described CD4BS bnAbs. Replacing V5 of JRFL with that of JRCSF had little effect on neutralization by sera 5394 and 5396 (**Supp. Table 1**). Likewise, a DT insertion in V5 of CH505.N197D, an insertion found in evolutionary variants of CH505 that reduces sensitivity to CD4BS bnAb CH103 (Liao et al., 2013), had a limited effect on most serum neutralization. However, the DT insertion did decrease neutralization of CH505.N197D by sera 5392 and 5397, by 2.4-fold and 5-fold, respectively, suggesting some nAbs elicited against the N197 glycan hole may also be interacting with V5.

## Discussion

Membrane Env is the sole target of HIV nAbs and hence is a reasonable platform for a vaccine, but its development has lagged far behind that of sEnv. Here, we describe MELs, which contain purified, mEnv spikes embedded and arrayed on the membrane of liposomes. VLP vaccines have been described previously but these typically have a low copy number of mEnv (Cantin et al., 2005). High Env VLPs (hVLPs) have been developed recently that display more than 100 spikes (Stano et al., 2017), but, like most VLPs, also carry Env debris, membrane proteins from the host cell, as well as viral proteins like capsid and matrix protein, which may divert the immune response. With MELs *any* Env can in principle be embedded, as GA can be used to fix mEnvs, the majority of which fail to form well-ordered sEnv trimers. Even reasonably stable mEnvs, such as ADA.CM and CH0505, can fail to form well-ordered sEnv trimers (Leaman & Zwick, 2013; Saunders et al., 2017), yet the latter were used here in MELs to elicit tier 2 nAbs.

Virosomes (Moser et al., 2007), nanodiscs (Witt et al., 2017), bicelles (Rantalainen et al., 2020), and ‘capture’ nanoparticles (Bale et al., 2017; Khairil Anuar et al., 2019; Leaman et al., 2015) are possible alternatives to MELs for displaying mEnv but either lack multivalent display or contain extraneous proteins that may divert the immune response. With MELs, T helper and B cell epitopes are exclusively from mEnv so may activate B cells efficiently *via* BCR crosslinking (Chackerian & Peabody, 2020). Size and composition of nanoparticles can affect trafficking in lymphatics as can antigen presentation and processing by dendritic cells (Brisse, Vrba, Kirk, Liang, & Ly, 2020); hence, future studies are warranted to determine whether changing MEL properties can be used to improve immune responses. Mixed MELs may also be prepared using many different Envs to potentially favor activation of B cells against conserved epitopes (Kanekiyo et al., 2019).

MELs were shown to elicit autologous nAbs in most animals but heterologous neutralization was limited. However, the sequential Env regimen used was likely suboptimal. CH505.N197D has a site-specific glycan hole at position 197 near the CD4BS and is sensitive to the bnAb CH103 UCA (*DPL and MBZ, ms. in preparation*). Neutralization titers to this mEnv rose rapidly on boosting and did so again on boosting with JRFL.TD15 that also lacks the N197 glycosylation site. It would be of interest to immunize rabbits and human Ig knock-in mice (Williams et al., 2017) using CH505.N197D and JRFL.TD15 in the prime and then boosting with glycan-restored Envs to see if CD4BS nAbs develop more effectively.

Among the MEL immune sera, antibody binding titers were low to the MPER and FP relative to the glycan deficient CD4BS or the variable loops, which are typically more immunogenic on Env (Crooks et al., 2015; Gray et al., 2011; Townsley, Li, Kozyrev, Cleveland, & Hu, 2016). However, significant sequence differences in the MPER and FP exist between the Envs we used, and weak binding titers were found to disappear on boosting. Hence, antibody responses might improve using Envs that have similar sequences in these regions, as well as high accessibility and/or affinity for bnAb UCAs (Zhang et al., 2019). Priming with epitope-specific peptides or scaffold proteins that engage bnAb UCAs, followed by MEL boosts, may also be tried. Stabilization of mEnv without chemical crosslinking and the use of specific lipids might enhance the immunogenicity of MELs as well. Finally, an MEL approach may be useful for eliciting antibodies against any membrane protein, pathogen or cancerous or diseased cells especially when membrane display influences key epitopes. Future work will focus on developing MEL vaccination in animals to more reproducibly elicit nAbs against the N197-proximal CD4BS, gp41 interface and MPER at the base of mEnv (Verkoczy et al., 2013; Xu et al., 2018).

## Materials and Methods

### Reagents

*(i) Cells*. TZM-bl cells were obtained from the NIH ARRRP. HEK 293T cells were purchased from the ATCC. High Env cell lines are discussed in more detail below.

*(ii) Antibodies*. Anti-HIV monoclonal antibodies b12, HGN194, 7B2, PGT128, PGT126, PGDM1400, VRC01, CH103, CH103 UCA, PGT151, 35O22, 10E8, F105, and 3BC176 were produced in-house as described previously (Gift, Leaman, Zhang, Kim, & Zwick, 2017). 2G12 and 2F5 were purchased from Polymun. Mouse antibody Chessie 8 was a gift from George Lewis (U. Maryland).

*(iii) Proteins and peptides*. Monomeric JRFL gp120 was purchased from Progenics, and JRFL gp41 MBP-fusion protein M41xt (aa 535-681) was produced in *E. coli* in-house (Nelson et al., 2007). The following peptides were synthesized by Genscript: JRV3 (^302^NTRLSIHIGPGRAFY TTGEIIGDI^327^), C34 (CH_3_O-^628^WMEWDREINNYTSLIHSLIEESQNQQEKNEQELL^663^-NH_2_), FP-bio (^512^AVGIGALFLGFLGAAGSTMGARS^534^-biotin), PID-bio (^592^LLGIWGCSQKLICTTA VPW^610^-biotin), MPER peptides PDT-081 (^654^EKNEQELLELDKWASLWNWFDITNWLW YIKKK^683^) and 94-1 (^671^NWFDITNWLWYIKKKK^683^), and CTT peptides CTT-2 (^725^RGPDRPEGIEEEGK^737^) and CTT-3 (^738^GERDRDRSIRL^748^).

### MEnv cell lines

The cell line ADA.CM (V4) has been described previously (Stano et al., 2017). The cell line CH505.N197D was similarly prepared using the CH505 Env sequence (Liao et al., 2013), in which a mutation N197D was introduced, by lentiviral transduction followed by iterative rounds using FACS to select for a PGT145^high^ b6^low^ phenotype. Likewise, the stable cell line expressing JRFL.TD15 mEnv was prepared using the sEnv sequence, described elsewhere (Guenaga et al., 2015), which was connected to the remaining MPER-TM-CTT sequence of wildtype JRFL. Of note, sequencing of DNA prepared from this cell line revealed a frame shift in the CTT at nucleotide position 2134 resulting in the following altered CTT sequence and early truncation: ^710^QGYSPCPSRPCCPPPAAPTAPRASRRRAASATATAPAAW*^749^. The cell lines were prepared fresh from freezer stocks, were passaged less than 20 times in DMEM containing 10% FBS, 2 mM L-glutamine, 100 U/ml penicillin, 100 μg/ml streptomycin and 2.5 μg/ml puromycin, and were kept from becoming overgrown by splitting every 3-4 d.

### Virus production

HIV was produced as a pseudotyped virus by transient transfection of HEK293T cells using Env plasmid DNA, pSG3ΔEnv backbone plasmid and 25 kDa PEI as the transfection reagent, as previously described (Agrawal et al., 2011).

### Production and purification of mEnv

Cell lines overexpressing mEnv were grown in shaker flasks at 37°C in 8% CO2 atmosphere to a density of 4 × 10^6^ cells/ml, then pelleted by centrifugation at 300 x *g* for 5 min. The cell pellet was resuspended in PBS and treated with 15 mM glutaraldehyde (GA; Sigma) at room temperature for 10 min. Unreacted GA was quenched for 15 min in 50 mM Tris-HCl, after which treated cells were solubilized for 20 min in 0.5% DDM (Sigma). Cell debris was removed by centrifugation at 3000 x *g* for 15 min at 4°C, then the supernatant was clarified by spinning at 80,000 x *g* in an Optima ultracentrifuge (Beckman) for 1 h at 4°C. MEnv was affinity purified using the trimer-specific bnAb PGT151 as previously described (Ringe et al., 2015; Sanders et al., 2013). Briefly, PGT151-coupled Protein A Sepharose beads (GE Healthcare) were added to the supernatant and rotated overnight at 4°C. The PGT151 beads were washed with TN75 (20 mM Tris, pH 8, 500 mM NaCl, 0.1% DDM, 0.003% sodium deoxycholate) and bound mEnv was eluted in 3M MgCl2. The trimer fraction was purified by size exclusion chromatography on a Superdex 200 10/300 GL column in TBS + 0.1% DDM + 0.003% sodium deoxycholate using an ÄKTA Pure 25L HPLC instrument (GE Healthcare) at a flow rate of 0.75 ml/min.

### Membrane Env liposome (MEL) production

Naked liposomes were prepared by dissolving POPC and cholesterol (Avanti) in chloroform at a 70:30 molar ratio. For liposomes to be analyzed by ELISA, 18:1 Biotinyl Cap PE (Avanti) was added to the lipids at a 2% molar ratio. Lipids were dried in a vacuum overnight, hydrated in PBS (20 mg/ml total lipid) with constant shaking for 2 h at 37°C, then sonicated for 30 s. The resulting liposomes were extruded 14 times through 1 μm, 0.8 μm, 0.4 μm, 0.2 μm, and 0.1 μm filters using a mini-extrusion device (Avanti) at room temperature (RT), then diluted to 4 mg/ml in PBS. DDM was added to a final concentration of 0.1% (1.9 mM, detergent:lipid molar ratio = 0.3:1) to destabilize the lipid bilayer (Lambert, Levy, Ranck, Leblanc, & Rigaud, 1998). Purified mEnv trimers were added at a 1:10 trimer:lipid ratio (w/w, and final concentrations of 1.9 mM DDM and 7.2 μM sodium deoxycholate) and the mixture was incubated at RT for 30 min. To remove detergent, polystyrene Bio-beads SM2 (Bio-Rad), pre-washed once with methanol and five times with water, were added at 40 mg/ml and the sample was rotated at RT for 30 min. Liposomes were treated thrice more with fresh Bio-beads at 4°C for 1 h, overnight, and 2 h, respectively. MELs in the supernatant were drawn off the polystyrene beads using a pipette and stored at 4°C.

### BN-PAGE

BN-PAGE was performed using the NativePAGE Gel System (ThermoFisher) as previously described (Leaman, Kinkead, & Zwick, 2010), except that samples were run on 3-12% gels. Coomassie staining was performed using Simply Blue Safe Stain (ThermoFisher) according to the manufacturer’s instructions.

### SDS-PAGE

Purified mEnv trimers (5 μg) were incubated in Laemmli Buffer (Bio-Rad) containing 50 mM dithiothreitol (DTT) for 5 min at 100°C before loading on 8-16% Tris-glycine gels (ThermoFisher). Gels were electrophoresed at RT in running buffer (25 mM Tris, 192 mM Glycine, 0.1% SDS, pH 8.3) at 150 V for 1 h, then Coomassie-stained as above.

### ELISAs

*(i) Capture ELISA. Galanthus nivalis* lectin (GNL) capture ELISAs were performed as described (Leaman et al., 2015). Briefly, microtiter wells were coated with GNL (5 μg/μl) in PBS overnight at 4°C, and then purified mEnv trimers (2 μg/μl) in PBS + 0.1% DDM were captured for 2 h at 37°C. Plates were blocked with 4% non-fat dry milk (NFDM) in PBS for 1 h at 37°C. Primary and HRP-conjugated secondary antibody incubations were performed for 1 h and 45 min, respectively, in PBS + 0.05% Tween-20 + 0.4% NFDM at 37°C.

*(ii) Direct ELISA*. Antigens (5 μg/μl) in PBS were coated onto microtiter wells overnight at 4°C, and ELISAs were performed as above without the antigen capture step.

*(iii) MEL ELISA*. MEL ELISAs were performed as above, except microtiter wells were coated with streptavidin (5 μg/μl) then blocked with 4% non-fat dry milk. MELs that incorporated biotinylated DOPE were captured on wells at 37°C for 2 h. Subsequent steps were done as above, but without detergent.

*(iv) Competition ELISA*. A capture ELISA was performed as above except serial dilutions of sera were added to wells at twice the final concentration. After 5 min, biotinylated bnAb was added at a concentration previously determined to produce 50% maximum signal and the mixture was incubated for 1 h at 37°C. BnAb binding was detected using streptavidin-HRP (Jackson).

### Nanoparticle Tracking Analysis (NTA)

A NanoSight N300 instrument (Malvern) was used to determine the dispersity and size of liposomes and MELs following a previously described method (Stano et al., 2017).

### Negative stain EM

Liposomes were applied for 2 min onto glow discharged, carbon-coated 400-Cu mesh grids (Electron Microscopy Sciences). Excess sample was removed by gentle contact with tissue paper, and the grids were placed on a droplet of 2% phosphotungstic acid (PTA) solution (pH 6.9) for 2 min. Excess stain was removed and grids were examined on a Philips CM100 electron microscope (FEI) at 80 kV. Images were acquired using a Megaview III charge-coupled device (CCD) camera (Olympus Soft Imaging Solutions).

### Rabbit immunization

Six New Zealand White (NZW) rabbits were immunized with a MEL formulation containing a total of 250 μg mEnv added to 100 μg CpG ODN 2007 (Invivogen) and Alum (Pierce) adjuvant, with a MEL:Alum ratio of 1:2 by volume. Injections were made bilaterally with 0.6 ml per injection *via* the subcutaneous route within 1 h of the formulation. MELs were used within 3 d of preparation. Blood was drawn from the marginal ear vein 4 d and 10 d post-injection for the preparation of PBMC and serum, respectively.

### Neutralization assays

Pseudotyped HIV-1 neutralization assays were performed using TZM-bl target cells, as previously described (Leaman et al., 2015). Briefly, TZM-bl cells were seeded onto a 96-well plate in 100 μL of growth media and incubated overnight at 37°C before the addition of virus. Virus was co-incubated with antibody or serum (previously heat-inactivated at 56°C for 30 min) at 37°C for 1 h and then added to TZM-bl cells. Infectivity was determined 72 h later by adding Bright-Glo (Promega) and measuring luciferase activity using a Synergy H1 plate reader (BioTek).

## Acknowledgments

We would like to thank Daniel Sands, Mark Ochoa and Trevor Biddle for technical support. We thank Jeong Hyun Lee, Jidnyasa Ingale, Shridhar Bale, and John Elder for helpful discussions on mEnv purification and MEL production. The Scripps Research Microscopy Core performed electron microscopy and Scripps Research Animal Resources carried out all animal procedures. Funding support was from the National Institutes of Health (NIAID) grants AI114401 and AI098602 (M.B.Z.), P01 AI104722 (R.T. Wyatt), as well as the James B. Pendleton Charitable Trust (M.B.Z.) to allow the purchase of an AKTA Pure instrument.

---

**Supplemental Table 1.**
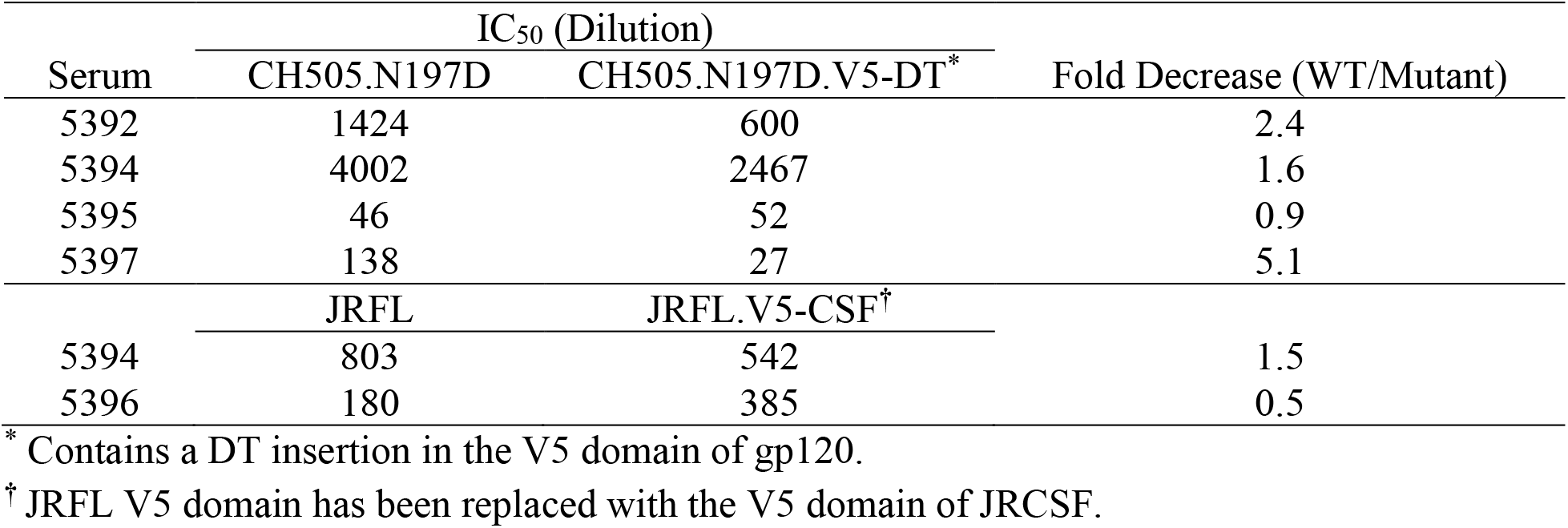
Effect of V5 substitutions on serum neutralization.

**Supp. Figure 1.**
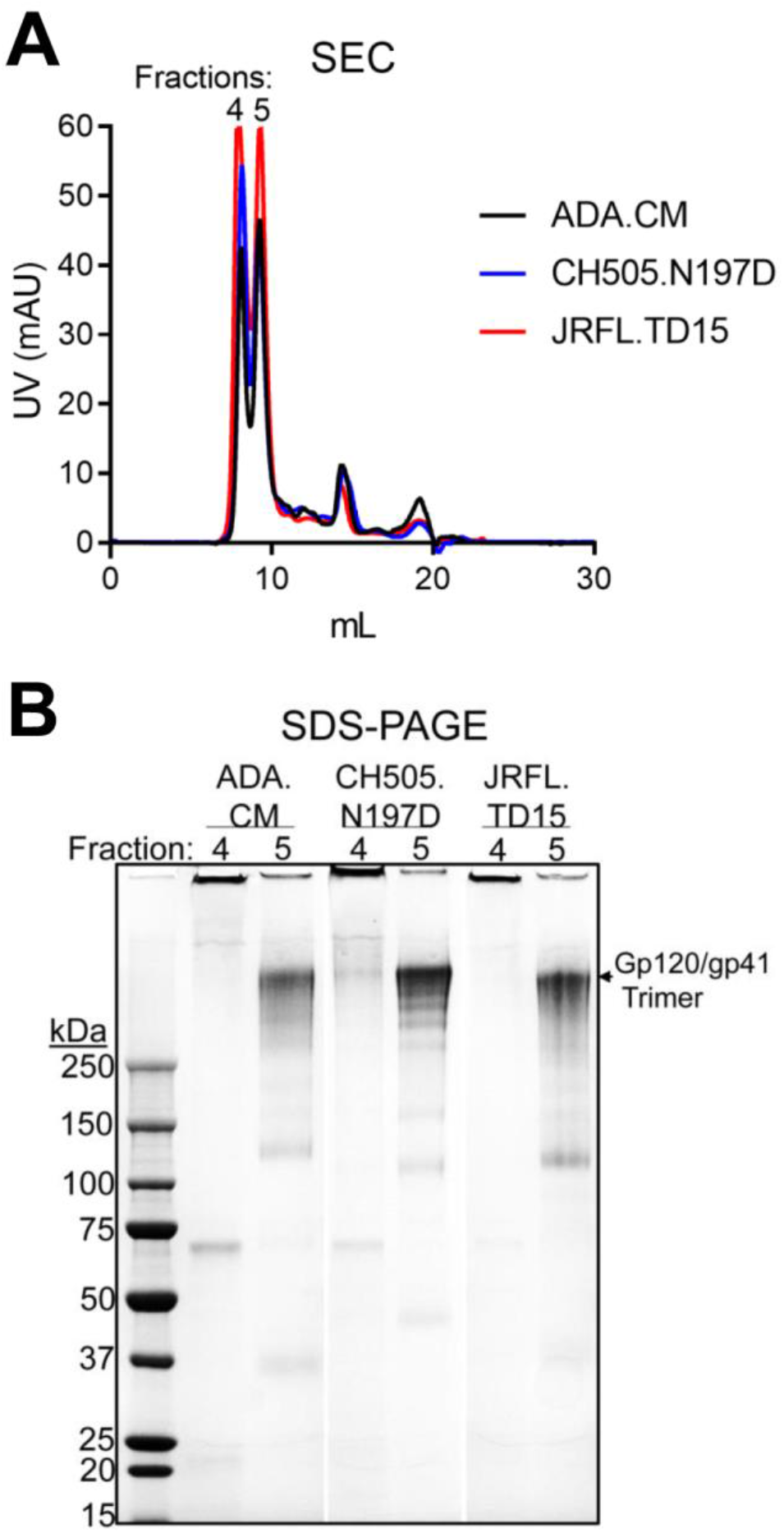
Separation of mEnv trimers from non-trimeric Env. (**A**) mEnv, GA crosslinked and PGT151 affinity-purified, was analyzed using size exclusion chromatography (SEC). (**B**) Two mEnv containing fractions from SEC were analyzed by denaturing Coomassie SDS-PAGE.

**Supp. Figure 2.**
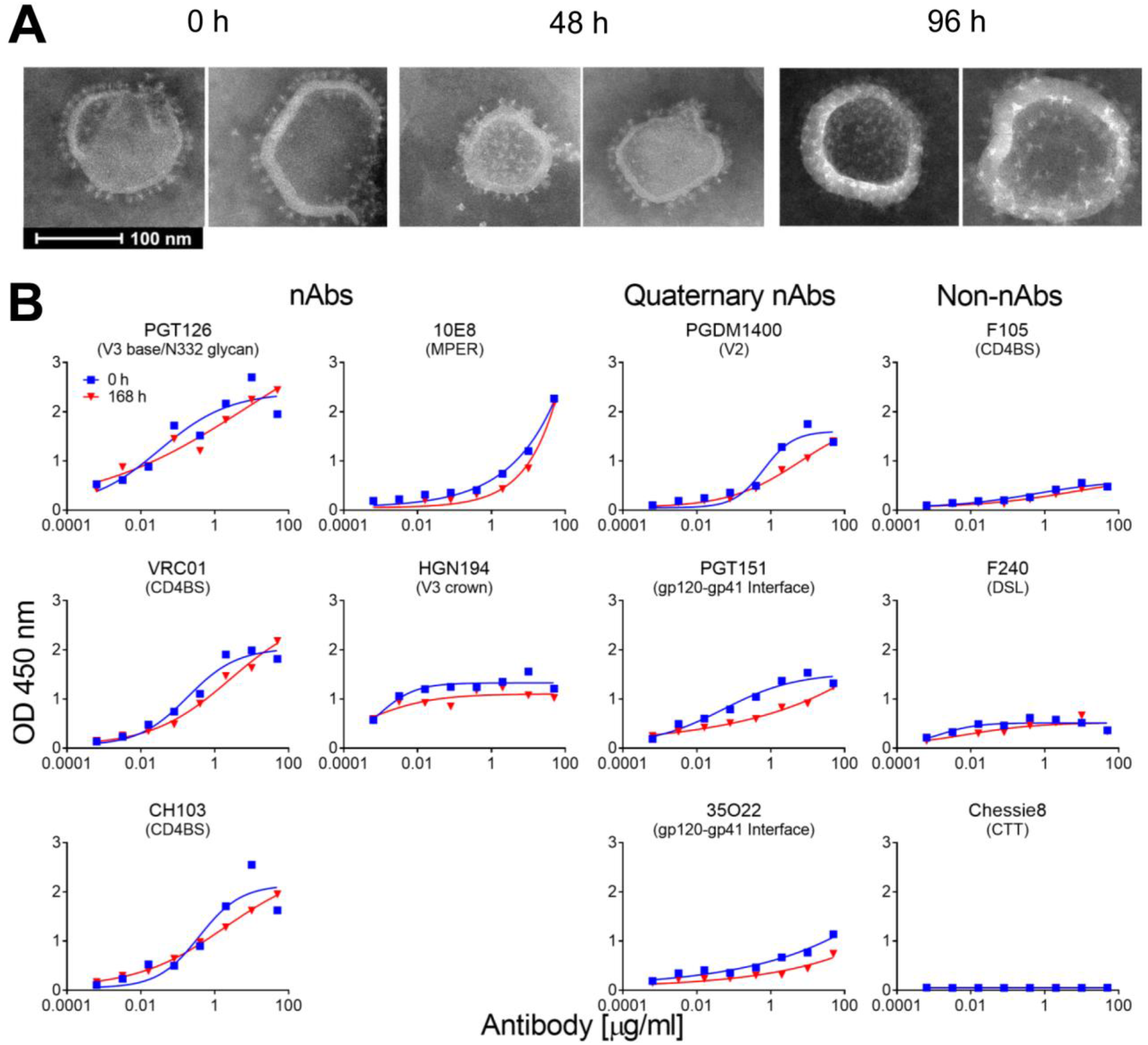
Stability of MELs in solution. (**A**) Negative stain EM images of MELs following 0, 48 and 96 h incubation periods at 37°C. (**B**) Antibody binding properties of MELs before and after incubation for 168 h at 37°C, as analyzed by streptavidin-capture ELISA as in Figure 2D.

**Supp. Figure 3.**
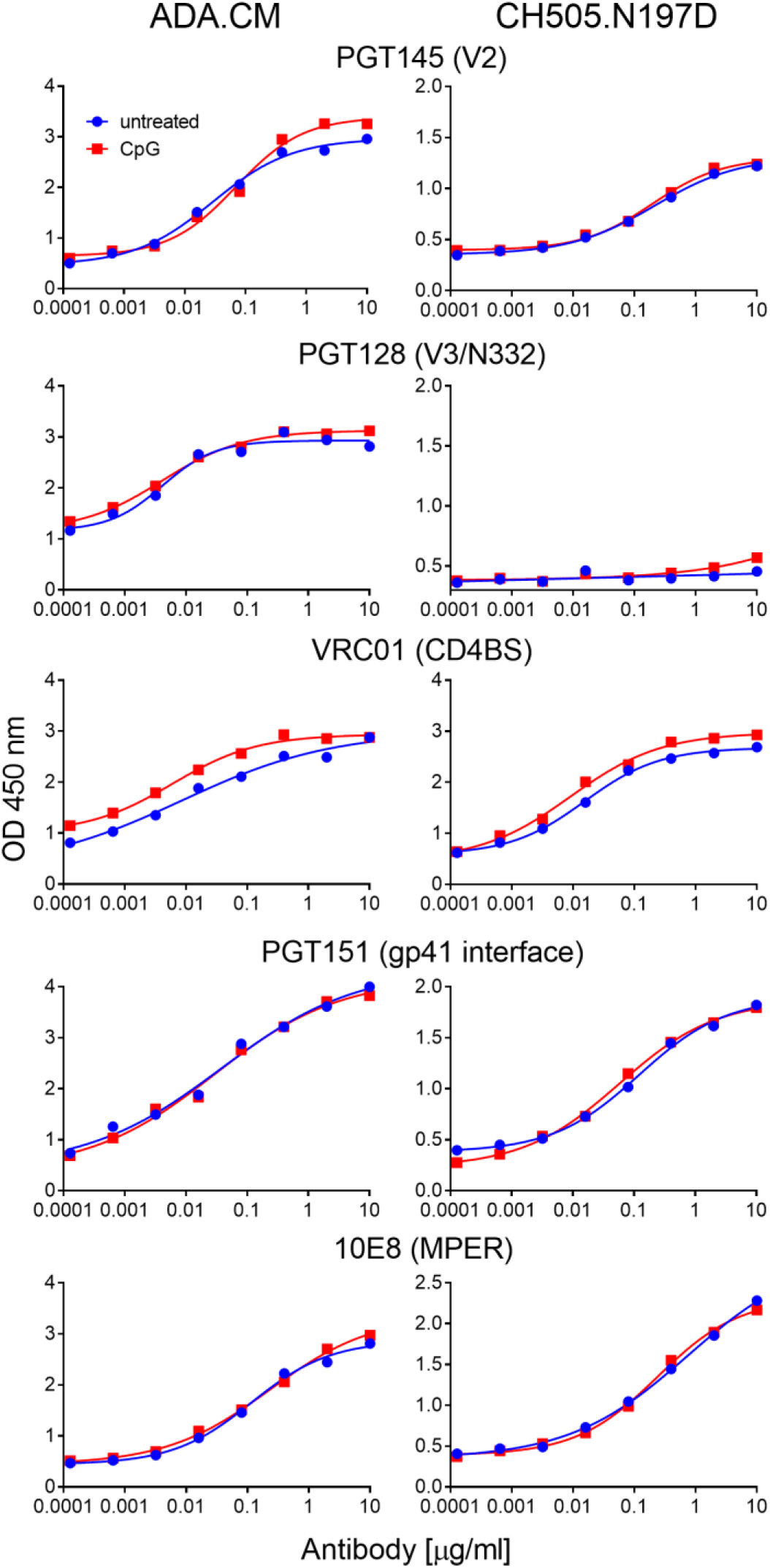
CpG ODN 2007 adjuvant does not occlude major bnAb epitopes. Antibody binding to immobilized ADA.CM and CH505.N197D trimers was determined by ELISA in the presence or absence of 15 μg/ml CpG ODN 2007.

**Supp. Figure 4.**
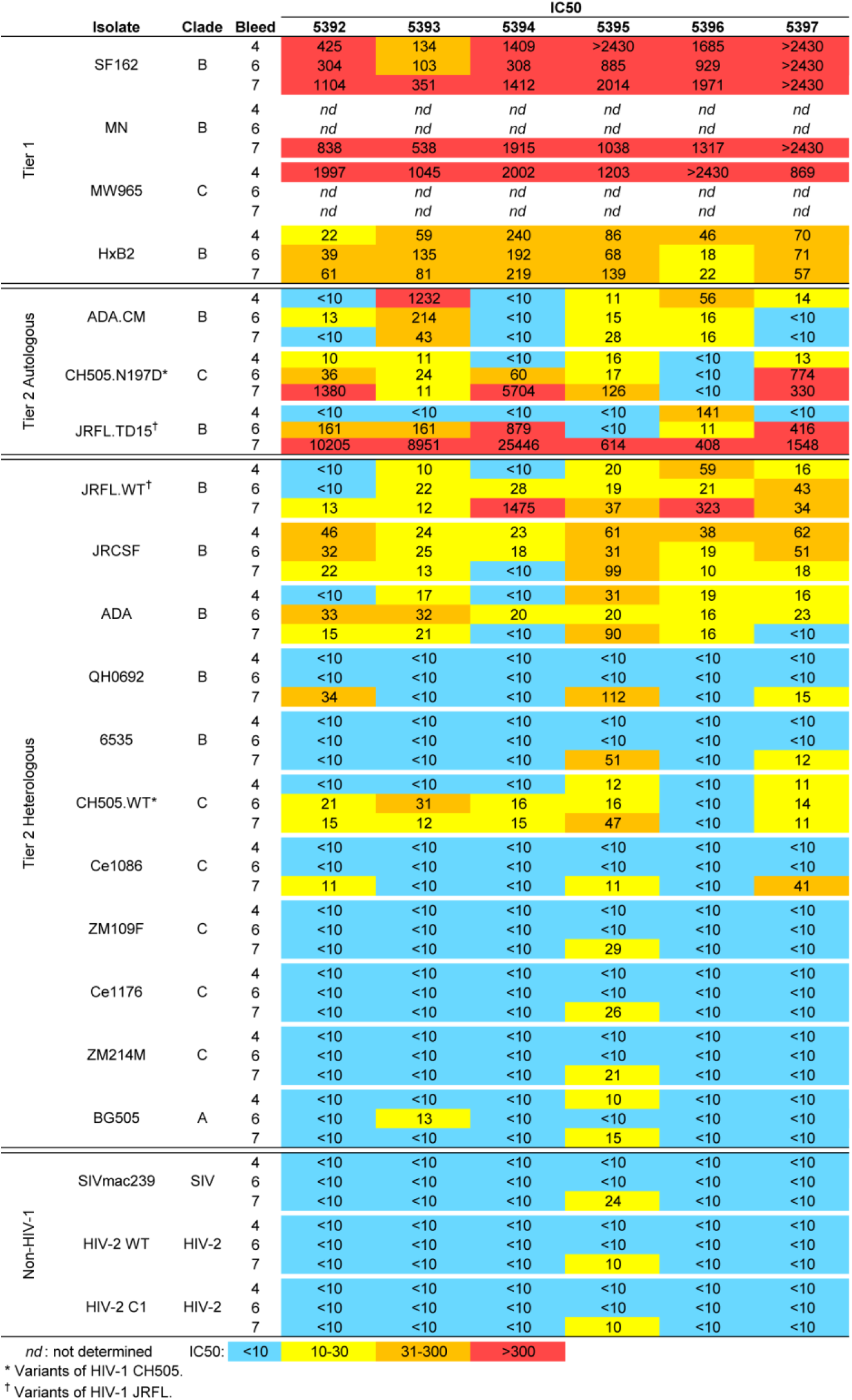
HIV neutralization breadth and potency of MEL immunized rabbit sera. Sera taken from rabbits were tested in neutralization assays against a cross-clade panel of autologous and heterologous HIV isolates. Data shown are reciprocal serum dilution at the IC_50_ from bleeds 4 (post ADA.CM), 6 (post CH505.N197D) and 7 (post JRFL.TD15) for each isolate.

**Supp. Figure 5.**
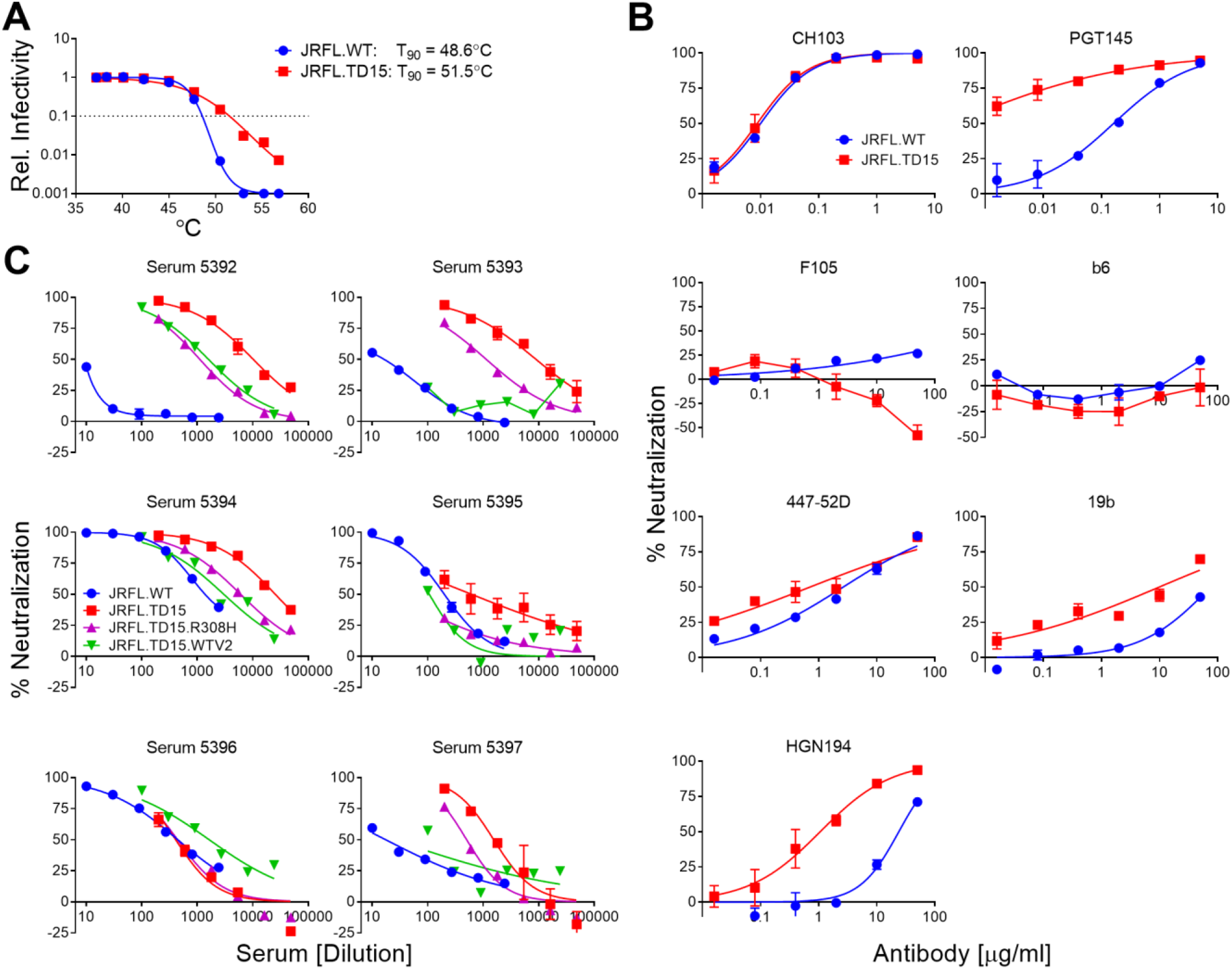
Stability and neutralization sensitivity of HIV JRFL.TD15. (**A**) Stability of function of JRFL.TD15 and JRFL.WT mEnvs was studied by determining the relative infectivity of cognate pseudovirions incubated for an hour at different temperatures; the temperature at which infectivity is reduced by 90% (T90) is indicated to the left. (**B**) Neutralization of JRFL.TD15 by narrow neutralizing antibodies against V3 (447-52D, 19b, and HGN194) and CD4BS (F105 and b6), as well as by bnAbs to the CD4BS (CH103) and to V2 (PGT145). (**C**) MEL-immunized rabbit sera were assayed for neutralization against JRFL.WT, JRFL.TD15 and mutants JRFL.TD15.R308H and JRFL.TD15.WTV2 that bear JRFL.WT V3 and V2 domains, respectively.

**Supp. Figure 6.**
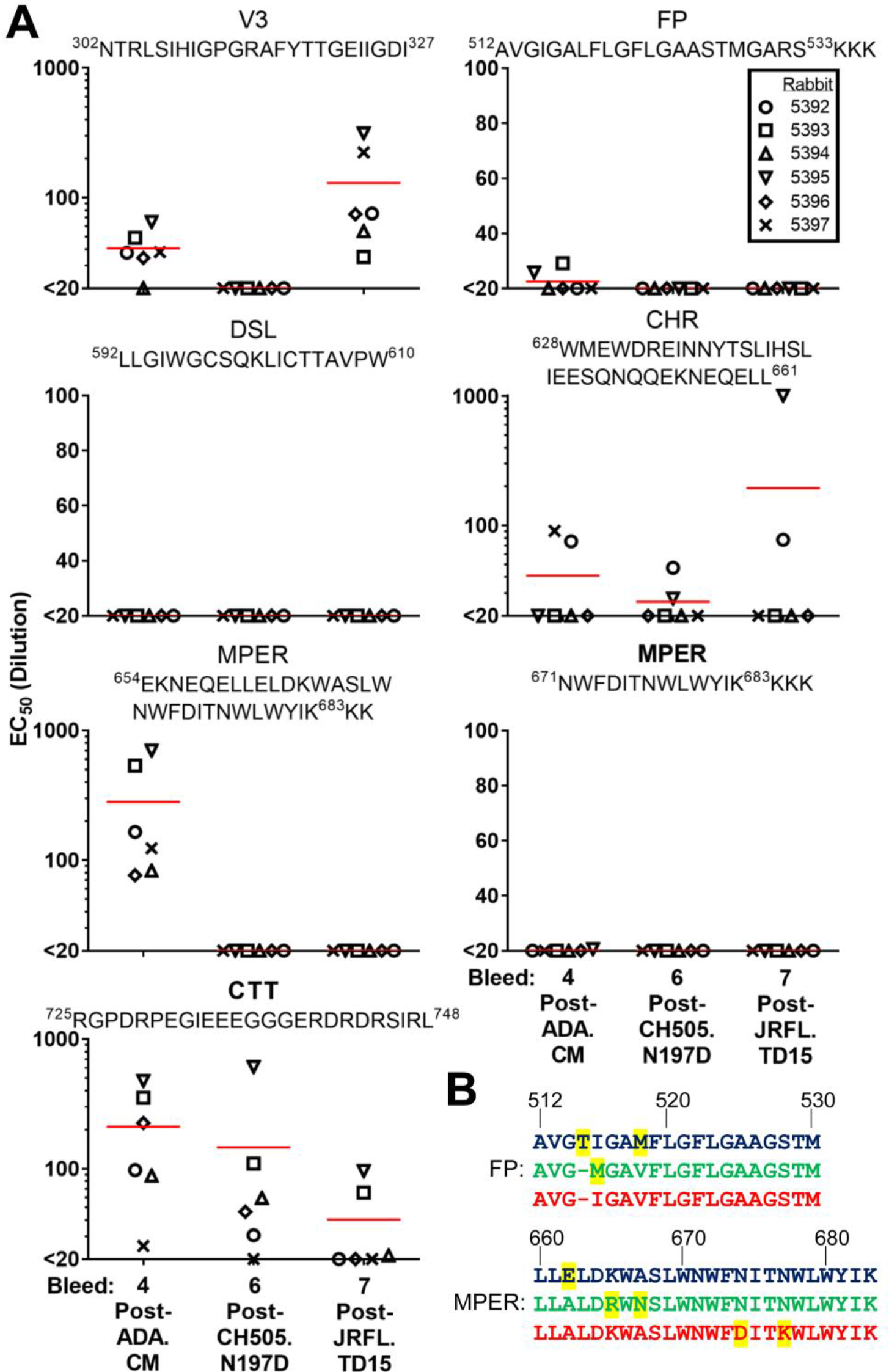
Binding specificities in sera of MEL immunized rabbits. (**A**) Sera from bleeds 4 (post-ADA.CM), 6 (post-CH505.N197D) and 7 (post-JRFL.TD15) were tested in an ELISA for the ability to bind various antigens and peptides of HIV Env. (**B**) Alignment of the FP (a.a. 512-530) and MPER (a.a. 660-683) of mEnvs used in the sequential immunization. ADA.CM sequence is in blue, CH505.N197D sequence is in green, and JRFL.TD15 sequence is in red. Residues that differ in one isolate relative to the other two are highlighted in yellow.

## References

Agrawal, N., Leaman, D. P., Rowcliffe, E., Kinkead, H., Nohria, R., Akagi, J., … Zwick, M. B. (2011). Functional Stability of Unliganded Envelope Glycoprotein Spikes among Isolates of Human Immunodeficiency Virus Type 1 (HIV-1). PLoS One, 6(6), e21339. doi:10.1371/journal.pone.0021339

Bale, S., Goebrecht, G., Stano, A., Wilson, R., Ota, T., Tran, K., … Wyatt, R. T. (2017). Covalent linkage of HIV-1 trimers to synthetic liposomes elicits improved B cell and antibody responses. J Virol. doi:10.1128/jvi.00443-17

Bjorkman, P. J. (2020). Can we use structural knowledge to design a protective vaccine against HIV-1? Hla, 95(2), 95–103. doi:10.1111/tan.13759

Brisse, M., Vrba, S. M., Kirk, N., Liang, Y., & Ly, H. (2020). Emerging Concepts and Technologies in Vaccine Development. Front Immunol, 11, 583077. doi:10.3389/fimmu.2020.583077

Cantin, R., Methot, S., & Tremblay, M. J. (2005). Plunder and stowaways: incorporation of cellular proteins by enveloped viruses. J Virol, 79(11), 6577–6587.

Chackerian, B., & Peabody, D. S. (2020). Factors That Govern the Induction of Long-Lived Antibody Responses. Viruses, 12(1). doi:10.3390/v12010074

Crooks, E. T., Moore, P. L., Franti, M., Cayanan, C. S., Zhu, P., Jiang, P., … Binley, J. M. (2007). A comparative immunogenicity study of HIV-1 virus-like particles bearing various forms of envelope proteins, particles bearing no envelope and soluble monomeric gp120. Virology, 366(2), 245–262. doi:10.1016/j.virol.2007.04.033

Crooks, E. T., Tong, T., Chakrabarti, B., Narayan, K., Georgiev, I. S., Menis, S., … Binley, J. M. (2015). Vaccine-Elicited Tier 2 HIV-1 Neutralizing Antibodies Bind to Quaternary Epitopes Involving Glycan-Deficient Patches Proximal to the CD4 Binding Site. PLoS Pathog, 11(5), e1004932. doi:10.1371/journal.ppat.1004932

Dubrovskaya, V., Tran, K., Ozorowski, G., Guenaga, J., Wilson, R., Bale, S., … Wyatt, R. T. (2019). Vaccination with Glycan-Modified HIV NFL Envelope Trimer-Liposomes Elicits Broadly Neutralizing Antibodies to Multiple Sites of Vulnerability. Immunity, 51(5), 915–929.e917. doi:10.1016/j.immuni.2019.10.008

Fera, D., Schmidt, A. G., Haynes, B. F., Gao, F., Liao, H. X., Kepler, T. B., & Harrison, S. C. (2014). Affinity maturation in an HIV broadly neutralizing B-cell lineage through reorientation of variable domains. Proc Natl Acad Sci U S A, 111(28), 10275–10280. doi:10.1073/pnas.1409954111

Geertsma, E. R., Nik Mahmood, N. A., Schuurman-Wolters, G. K., & Poolman, B. (2008). Membrane reconstitution of ABC transporters and assays of translocator function. Nat Protoc, 3(2), 256–266. doi:10.1038/nprot.2007.519

Gift, S. K., Leaman, D. P., Zhang, L., Kim, A. S., & Zwick, M. B. (2017). Functional Stability of HIV-1 Envelope Trimer Affects Accessibility to Broadly Neutralizing Antibodies at Its Apex. J Virol, 91(24). doi:10.1128/jvi.01216-17

Gilbert, P., Wang, M., Wrin, T., Petropoulos, C., Gurwith, M., Sinangil, F., … Montefiori, D. C. (2010). Magnitude and breadth of a nonprotective neutralizing antibody response in an efficacy trial of a candidate HIV-1 gp120 vaccine. J Infect Dis, 202(4), 595–605. doi:10.1086/654816

Gray, E. S., Madiga, M. C., Hermanus, T., Moore, P. L., Wibmer, C. K., Tumba, N. L., … Morris, L. (2011). The neutralization breadth of HIV-1 develops incrementally over four years and is associated with CD4+ T cell decline and high viral load during acute infection. J Virol, 85(10), 4828–4840. doi:10.1128/jvi.00198-11

Guenaga, J., Dubrovskaya, V., de Val, N., Sharma, S. K., Carrette, B., Ward, A. B., & Wyatt, R. T. (2015). Structure-Guided Redesign Increases the Propensity of HIV Env To Generate Highly Stable Soluble Trimers. J Virol, 90(6), 2806–2817. doi:10.1128/jvi.02652-15

Haynes, B. F., Burton, D. R., & Mascola, J. R. (2019). Multiple roles for HIV broadly neutralizing antibodies. Sci Transl Med, 11(516). doi:10.1126/scitranslmed.aaz2686

Hessell, A. J., Malherbe, D. C., Pissani, F., McBurney, S., Krebs, S. J., Gomes, M., … Haigwood, N. L. (2016). Achieving Potent Autologous Neutralizing Antibody Responses against Tier 2 HIV-1 Viruses by Strategic Selection of Envelope Immunogens. J Immunol, 196(7), 3064–3078. doi:10.4049/jimmunol.1500527

Hu, J. K., Crampton, J. C., Cupo, A., Ketas, T., van Gils, M. J., Sliepen, K., … Crotty, S. (2015). Murine Antibody Responses to Cleaved Soluble HIV-1 Envelope Trimers Are Highly Restricted in Specificity. J Virol, 89(20), 10383–10398. doi:10.1128/jvi.01653-15

Julien, J. P., Cupo, A., Sok, D., Stanfield, R. L., Lyumkis, D., Deller, M. C., … Wilson, I. A. (2013). Crystal structure of a soluble cleaved HIV-1 envelope trimer. Science, 342(6165), 1477–1483. doi:10.1126/science.1245625

Kanekiyo, M., Joyce, M. G., Gillespie, R. A., Gallagher, J. R., Andrews, S. F., Yassine, H. M., … Graham, B. S. (2019). Mosaic nanoparticle display of diverse influenza virus hemagglutinins elicits broad B cell responses. Nat Immunol, 20(3), 362–372. doi:10.1038/s41590-018-0305-x

Khairil Anuar, I. N. A., Banerjee, A., Keeble, A. H., Carella, A., Nikov, G. I., & Howarth, M. (2019). Spy&Go purification of SpyTag-proteins using pseudo-SpyCatcher to access an oligomerization toolbox. Nat Commun, 10(1), 1734. doi:10.1038/s41467-019-09678-w

Kong, L., He, L., de Val, N., Vora, N., Morris, C. D., Azadnia, P., … Zhu, J. (2016). Uncleaved prefusion-optimized gp140 trimers derived from analysis of HIV-1 envelope metastability. Nat Commun, 7, 12040. doi:10.1038/ncomms12040

Krebs, S. J., Kwon, Y. D., Schramm, C. A., Law, W. H., Donofrio, G., Zhou, K. H., … Doria-Rose, N. A. (2019). Longitudinal Analysis Reveals Early Development of Three MPER-Directed Neutralizing Antibody Lineages from an HIV-1-Infected Individual. Immunity, 50(3), 677–691.e613. doi:10.1016/j.immuni.2019.02.008

Lambert, O., Levy, D., Ranck, J. L., Leblanc, G., & Rigaud, J. L. (1998). A new “gel-like” phase in dodecyl maltoside-lipid mixtures: implications in solubilization and reconstitution studies. Biophys J, 74(2 Pt 1), 918–930. doi:10.1016/s0006-3495(98)74015-9

Leaman, D. P., Kinkead, H., & Zwick, M. B. (2010). In-solution virus capture assay helps deconstruct heterogeneous antibody recognition of human immunodeficiency virus type 1. J Virol, 84(7), 3382–3395.

Leaman, D. P., Lee, J. H., Ward, A. B., & Zwick, M. B. (2015). Immunogenic Display of Purified Chemically Cross-Linked HIV-1 Spikes. J Virol, 89(13), 6725–6745. doi:10.1128/jvi.03738-14

Leaman, D. P., & Zwick, M. B. (2013). Increased functional stability and homogeneity of viral envelope spikes through directed evolution. PLoS Pathog, 9(2), e1003184. doi:10.1371/journal.ppat.1003184

Lee, J. H., Ozorowski, G., & Ward, A. B. (2016). Cryo-EM structure of a native, fully glycosylated, cleaved HIV-1 envelope trimer. Science, 351(6277’), 1043–1048. doi:10.1126/science.aad2450

Liao, H. X., Lynch, R., Zhou, T., Gao, F., Alam, S. M., Boyd, S. D., … Haynes, B. F. (2013). Co-evolution of a broadly neutralizing HIV-1 antibody and founder virus. Nature, 496(7446), 469–476. doi:10.1038/nature12053

Lu, M., Ma, X., Castillo-Menendez, L. R., Gorman, J., Alsahafi, N., Ermel, U., … Mothes, W. (2019). Associating HIV-1 envelope glycoprotein structures with states on the virus observed by smFRET. Nature, 568(7752), 415–419. doi:10.1038/s41586-019-1101-y

Mascola, J. R., Snyder, S. W., Weislow, O. S., Belay, S. M., Belshe, R. B., Schwartz, D. H., … Burke, D. S. (1996). Immunization with envelope subunit vaccine products elicits neutralizing antibodies against laboratory-adapted but not primary isolates of human immunodeficiency virus type 1. J Infect Dis, 173(2), 340–348.

McCoy, L. E., van Gils, M. J., Ozorowski, G., Messmer, T., Briney, B., Voss, J. E., … Burton, D. R. (2016). Holes in the Glycan Shield of the Native HIV Envelope Are a Target of Trimer-Elicited Neutralizing Antibodies. Cell Rep, 16(9), 2327–2338. doi:10.1016/j.celrep.2016.07.074

McGuire, A. T., Gray, M. D., Dosenovic, P., Gitlin, A. D., Freund, N. T., Petersen, J., … Stamatatos, L. (2016). Specifically modified Env immunogens activate B-cell precursors of broadly neutralizing HIV-1 antibodies in transgenic mice. Nat Commun, 7, 10618. doi:10.1038/ncomms10618

Morris, C. D., Azadnia, P., de Val, N., Vora, N., Honda, A., Giang, E., … Zhu, J. (2017). Differential Antibody Responses to Conserved HIV-1 Neutralizing Epitopes in the Context of Multivalent Scaffolds and Native-Like gp140 Trimers. MBio(1). doi:10.1128/mBio.00036-17

Moser, C., Amacker, M., Kammer, A. R., Rasi, S., Westerfeld, N., & Zurbriggen, R. (2007). Influenza virosomes as a combined vaccine carrier and adjuvant system for prophylactic and therapeutic immunizations. Expert Rev Vaccines, 6(5), 711–721. doi:10.1586/14760584.6.5.711

Nelson, J. D., Brunel, F. M., Jensen, R., Crooks, E. T., Cardoso, R. M., Wang, M., … Zwick, M. B. (2007). An affinity-enhanced neutralizing antibody against the membrane-proximal external region of human immunodeficiency virus type 1 gp41 recognizes an epitope between those of 2F5 and 4E10. J Virol, 81(8), 4033–4043.

Ozorowski, G., Cupo, A., Golabek, M., LoPiccolo, M., Ketas, T. A., Cavallary, M., … Moore, J. P. (2018). Effects of Adjuvants on HIV-1 Envelope Glycoprotein SOSIP Trimers In Vitro. J Virol, 92(13). doi:10.1128/jvi.00381-18

Pardi, N., Hogan, M. J., Porter, F. W., & Weissman, D. (2018). mRNA vaccines - a new era in vaccinology. Nat Rev Drug Discov, 17(4), 261–279. doi:10.1038/nrd.2017.243

Poon, B., Hsu, J. F., Gudeman, V., Chen, I. S., & Grovit-Ferbas, K. (2005). Formaldehyde-treated, heat-inactivated virions with increased human immunodeficiency virus type 1 env can be used to induce high-titer neutralizing antibody responses. J Virol, 79(16), 10210–10217. doi:10.1128/jvi.79.16.10210-10217.2005

Rantalainen, K., Berndsen, Z. T., Antanasijevic, A., Schiffner, T., Zhang, X., Lee, W. H., … Ward, A. B. (2020). HIV-1 Envelope and MPER Antibody Structures in Lipid Assemblies. Cell Rep, 31(4), 107583. doi:10.1016/j.celrep.2020.107583

Rerks-Ngarm, S., Pitisuttithum, P., Nitayaphan, S., Kaewkungwal, J., Chiu, J., Paris, R., … Kim, J. H. (2009). Vaccination with ALVAC and AIDSVAX to prevent HIV-1 infection in Thailand. N Engl J Med, 361(23), 2209–2220. doi:10.1056/NEJMoa0908492

Rigaud, J. L., Mosser, G., Lacapere, J. J., Olofsson, A., Levy, D., & Ranck, J. L. (1997). Bio-Beads: an efficient strategy for two-dimensional crystallization of membrane proteins. J Struct Biol, 118(3), 226–235. doi:10.1006/jsbi.1997.3848

Ringe, R. P., Sanders, R. W., Yasmeen, A., Kim, H. J., Lee, J. H., Cupo, A., … Moore, J. P. (2013). Cleavage strongly influences whether soluble HIV-1 envelope glycoprotein trimers adopt a native-like conformation. Proc Natl Acad Sci U S A, 110(45), 18256–18261. doi:10.1073/pnas.1314351110

Ringe, R. P., Yasmeen, A., Ozorowski, G., Go, E. P., Pritchard, L. K., Guttman, M., … Moore, J. P. (2015). Influences on the Design and Purification of Soluble, Recombinant Native-Like HIV-1 Envelope Glycoprotein Trimers. J Virol, 89(23), 12189–12210. doi:10.1128/jvi.01768-15

Sanders, R. W., Derking, R., Cupo, A., Julien, J. P., Yasmeen, A., de Val, N., … Moore, J. P. (2013). A next-generation cleaved, soluble HIV-1 Env trimer, BG505 SOSIP.664 gp140, expresses multiple epitopes for broadly neutralizing but not non-neutralizing antibodies. PLoS Pathog, 9(9), e1003618. doi:10.1371/journal.ppat.1003618

Sanders, R. W., van Gils, M. J., Derking, R., Sok, D., Ketas, T. J., Burger, J. A., … Moore, J. P. (2015). HIV-1 VACCINES. HIV-1 neutralizing antibodies induced by native-like envelope trimers. Science, 349(6244), aac4223. doi:10.1126/science.aac4223

Sanders, R. W., Vesanen, M., Schuelke, N., Master, A., Schiffner, L., Kalyanaraman, R., … Moore, J. P. (2002). Stabilization of the soluble, cleaved, trimeric form of the envelope glycoprotein complex of human immunodeficiency virus type 1. J Virol, 76(17), 8875–8889.

Saunders, K. O., Verkoczy, L. K., Jiang, C., Zhang, J., Parks, R., Chen, H., … Haynes, B. F. (2017). Vaccine Induction of Heterologous Tier 2 HIV-1 Neutralizing Antibodies in Animal Models. Cell Rep, 21(13), 3681–3690. doi:10.1016/j.celrep.2017.12.028

Schiffner, T., Pallesen, J., Russell, R. A., Dodd, J., de Val, N., LaBranche, C. C., … Sattentau, Q. J. (2018). Structural and immunologic correlates of chemically stabilized HIV-1 envelope glycoproteins. PLoS Pathog, 14(5), e1006986. doi:10.1371/journal.ppat.1006986

Schommers, P., Gruell, H., Abernathy, M. E., Tran, M. K., Dingens, A. S., Gristick, H. B., … Klein, F. (2020). Restriction of HIV-1 Escape by a Highly Broad and Potent Neutralizing Antibody. Cell, 180(3), 471–489.e422. doi:10.1016/j.cell.2020.01.010

Seddon, A. M., Curnow, P., & Booth, P. J. (2004). Membrane proteins, lipids and detergents: not just a soap opera. Biochim Biophys Acta, 1666(1-2), 105–117. doi:10.1016/j.bbamem.2004.04.011

Sharma, S. K., de Val, N., Bale, S., Guenaga, J., Tran, K., Feng, Y., … Wyatt, R. T. (2015). Cleavage-independent HIV-1 Env trimers engineered as soluble native spike mimetics for vaccine design. Cell Rep, 11(4), 539–550. doi:10.1016/j.celrep.2015.03.047

Soldemo, M., Adori, M., Stark, J. M., Feng, Y., Tran, K., Wilson, R., … Karlsson Hedestam, G. B. (2017). Glutaraldehyde Cross-linking of HIV-1 Env Trimers Skews the Antibody Subclass Response in Mice. Front Immunol, 8, 1654. doi:10.3389/fimmu.2017.01654

Spearman, P. (2006). Current progress in the development of HIV vaccines. Curr Pharm Des, 12(9), 1147–1167.

Spearman, P., Lally, M. A., Elizaga, M., Montefiori, D., Tomaras, G. D., McElrath, M. J., … Corey, L. J. (2011). A trimeric, V2-deleted HIV-1 envelope glycoprotein vaccine elicits potent neutralizing antibodies but limited breadth of neutralization in human volunteers. J Infect Dis, 203(8), 1165–1173. doi:10.1093/infdis/jiq175

Stadtmueller, B. M., Bridges, M. D., Dam, K. M., Lerch, M. T., Huey-Tubman, K. E., Hubbell, W. L., & Bjorkman, P. J. (2018). DEER Spectroscopy Measurements Reveal Multiple Conformations of HIV-1 SOSIP Envelopes that Show Similarities with Envelopes on Native Virions. Immunity, 49(2), 235–246.e234. doi:10.1016/j.immuni.2018.06.017

Stano, A., Leaman, D. P., Kim, A. S., Zhang, L., Autin, L., Ingale, J., … Zwick, M. B. (2017). Dense array of spikes on HIV-1 virion particles. J Virol, 91(14), 415–417. doi:10.1128/jvi.00415-17

Steckbeck, J. D., Sun, C., Sturgeon, T. J., & Montelaro, R. C. (2013). Detailed topology mapping reveals substantial exposure of the “cytoplasmic” C-terminal tail (CTT) sequences in HIV-1 Env proteins at the cell surface. PLoS One, 8(5), e65220. doi:10.1371/journal.pone.0065220

Tomaras, G. D., & Plotkin, S. A. (2017). Complex immune correlates of protection in HIV-1 vaccine efficacy trials. Immunol Rev, 275(1), 245–261. doi:10.1111/imr.12514

Townsley, S., Li, Y., Kozyrev, Y., Cleveland, B., & Hu, S. L. (2016). Conserved Role of an N-Linked Glycan on the Surface Antigen of Human Immunodeficiency Virus Type 1 Modulating Virus Sensitivity to Broadly Neutralizing Antibodies against the Receptor and Coreceptor Binding Sites. J Virol, 90(2), 829–841. doi:10.1128/jvi.02321-15

Umotoy, J., Bagaya, B. S., Joyce, C., Schiffner, T., Menis, S., Saye-Francisco, K. L., … Landais, E. (2019). Rapid and Focused Maturation of a VRC01-Class HIV Broadly Neutralizing Antibody Lineage Involves Both Binding and Accommodation of the N276-Glycan. Immunity, 51(1), 141–154.e146. doi:10.1016/j.immuni.2019.06.004

UNAIDS. (2020). UNAIDS Data 2020. Retrieved from https://www.unaids.org/sites/default/files/media_asset/2020_aids-data-book_en.pdf

Verkoczy, L., Chen, Y., Zhang, J., Bouton-Verville, H., Newman, A., Lockwood, B., … Haynes, B. F. (2013). Induction of HIV-1 broad neutralizing antibodies in 2F5 knock-in mice: selection against membrane proximal external region-associated autoreactivity limits T-dependent responses. J Immunol, 191(5), 2538–2550. doi:10.4049/jimmunol.1300971

Voss, J. E., Andrabi, R., McCoy, L. E., de Val, N., Fuller, R. P., Messmer, T., … Burton, D. R. (2017). Elicitation of Neutralizing Antibodies Targeting the V2 Apex of the HIV Envelope Trimer in a Wild-Type Animal Model. Cell Rep, 21(1), 222–235. doi:10.1016/j.celrep.2017.09.024

Walker, L. M., Phogat, S. K., Chan-Hui, P. Y., Wagner, D., Phung, P., Goss, J. L., … Burton, D. R. (2009). Broad and potent neutralizing antibodies from an African donor reveal a new HIV-1 vaccine target. Science, 326(5950), 285–289.

Williams, W. B., Zhang, J., Jiang, C., Nicely, N. I., Fera, D., Luo, K., … Verkoczy, L. (2017). Initiation of HIV neutralizing B cell lineages with sequential envelope immunizations. Nat Commun, 8(1), 1732. doi:10.1038/s41467-017-01336-3

Witt, K. C., Castillo-Menendez, L., Ding, H., Espy, N., Zhang, S., Kappes, J. C., & Sodroski, J. (2017). Antigenic characterization of the human immunodeficiency virus (HIV-1) envelope glycoprotein precursor incorporated into nanodiscs. PLoS One, 12(2), e0170672. doi:10.1371/journal.pone.0170672

Xu, K., Acharya, P., Kong, R., Cheng, C., Chuang, G. Y., Liu, K., … Kwong, P. D. (2018). Epitope-based vaccine design yields fusion peptide-directed antibodies that neutralize diverse strains of HIV-1. Nat Med, 24(6), 857–867. doi:10.1038/s41591-018-0042-6

Yang, X., Wyatt, R., & Sodroski, J. (2001). Improved elicitation of neutralizing antibodies against primary human immunodeficiency viruses by soluble stabilized envelope glycoprotein trimers. J Virol, 75(3), 1165–1171. doi:10.1128/jvi.75.3.1165-1171.2001

Zhang, L., Irimia, A., He, L., Landais, E., Rantalainen, K., Leaman, D. P., … Zwick, M. B. (2019). An MPER antibody neutralizes HIV-1 using germline features shared among donors. Nat Commun, 10(1), 5389. doi:10.1038/s41467-019-12973-1

Zhou, T., Doria-Rose, N. A., Cheng, C., Stewart-Jones, G. B. E., Chuang, G. Y., Chambers, M., … Kwong, P. D. (2017). Quantification of the Impact of the HIV-1-Glycan Shield on Antibody Elicitation. Cell Rep, 19(4), 719–732. doi:10.1016/j.celrep.2017.04.013

